# Folded alpha helical putative new proteins from *Apilactobacillus kunkeei*

**DOI:** 10.1101/2023.08.08.552426

**Authors:** Weihua Ye, Phani Rama Krishna Behra, Karl Dyrhage, Christian Seeger, Joe D. Joiner, Elin Karlsson, Eva Andersson, Celestine N. Chi, Siv G. E. Andersson, Per Jemth

## Abstract

The emergence of new proteins is a central question in biology. Most tertiary protein folds known to date appear to have an ancient origin, but it is clear from bioinformatic analyses that new proteins continuously emerge in all organismal groups. However, there is a paucity of experimental data on new proteins regarding their structure and biophysical properties. We performed a detailed phylogenetic analysis and identified 48 putative open reading frames in the honeybee-associated bacterium *Apilactobacillus kunkeei* for which no homologs could be identified in closely-related species, suggesting that they could be relatively new on an evolutionary time scale and represent recently evolved proteins. Using circular dichroism-, fluorescence- and nuclear magnetic resonance spectroscopy we investigated five of these proteins and show that they are not intrinsically disordered, but populate alpha-helical dominated folded states with relatively low thermodynamic stability (0-3 kcal/mol). The data demonstrate that small new proteins readily adopt simple folded conformations suggesting that more complex tertiary structures can be continuously re-invented during evolution by fusion of such simple secondary structure elements. These findings have implications for the general view on protein evolution, where *de novo* emergence of folded proteins may be a common event.

## INTRODUCTION

New genes arise by duplication and divergence ^1^, exon rearrangements and gene fusion/fission events in which protein domains encoded by already existing genes are reused for new or modified functions ^2, 3^. Expansions of functional repertoires by duplication and domain shuffling events are commonly observed in protein families involved in signal transduction pathways as well as in gene regulation and transport systems. More recently, it has been shown that new genes can arise *de novo* from non-coding sequences ^4–6^, as first demonstrated for genes involved in male reproduction in fruit flies ^7, 8^. While the majority of genes encoding new proteins are lost because they do not confer a fitness advantage to the organism, some of the new proteins may be subject to positive selection and their genes retained in the genome of future generations. Furthermore, evolutionary experiments have demonstrated that randomly synthesized short open reading frames can confer new function, for example antibiotic resistance ^9,10^ or rescue of auxotrophy ^11^ in *Escherichia coli*.

However, despite these examples of gene birth from non-coding DNA, the mechanisms and frequencies with which new protein-coding genes are generated *de novo* and in particular the structure and function of the new proteins remain largely unknown. It is widely accepted that the origin of most of the recognized protein folds in present-day organisms are ancient, and were present in the last universal common ancestor ^12–14^. However, there is a growing body of data showing that new proteins constantly emerge in living organisms ^15^. It is conceptually very important to understand whether *de novo* emergence of protein domains and structural convergence ^16, 17^ are common or rare events. A key question is whether *de novo* proteins can fold into well-defined structures and converge on existing protein folds, or if new proteins are predominately intrinsically disordered as suggested by predictions ^18^.

We address this question by structural characterization of putative small new proteins expressed from short open reading frames (smORFs) in *Apilactobacillus kunkeii*, a defensive symbiont of honeybees that is highly abundant in the honey crop and the honeybee food products ^19–24^. The *A. kunkeii* isolates have genome sizes ranging from 1.49 - 1.64 Mb (mean 1.57 Mb), and gene contents ranging from 1345 -1504 genes (mean 1430) ^25^. A study of 104 closed genomes indicated that the population has an open pan-genome structure, meaning that the number of new chromosomal genes increases with the addition of every genome ^25^. However, the mechanism whereby these new genes have originated is not known, nor is it known if the new genes are temporary residents in the genome or code for proteins that confer new functions to the bacterial cell.

We have here investigated in detail a subset of *A. kunkeii* smORFs from the perspective of protein structure. We find that these small and potentially new proteins are not intrinsically disordered but instead adopt simple alpha-helical folds. Further evolution of such small structured proteins, for example by fusion of their genes, could generate larger folded proteins with implications for the general thinking about emergence of novel protein folds.

## RESULTS

### Identification of recently evolved smORFs coding for hypothetical proteins

The starting point for this analysis was the 1466 predicted genes in the genome of *A. kunkeei* strain A0901. A BLASTP search against the NR database ver 2023-01-10 showed that 137 genes had less than 100 hits to species outside *A. kunkeei* (with the NCBI-taxonomy ids “148814”, “1419324”, “1423768” and E less than 1e-03) (<u>Figure S1a</u>). Of these, 48 open reading frames with a minimum length of 30 bp and an average length of ∼111 bp showed no hits or only a few (max five) hits to hypothetical genes in bacteria with taxonomic names other than *A. kunkeei* (<u>Table S1, Figure S1b</u>). We calculated average nucleotide sequence identity values (ANI) for all pairwise comparisons of these taxa to the *A. kunkeei* isolates (<u>Table S2</u>), and used the values as a proxy for evolutionary relatedness. Most taxa showed ANI values above 97% and should be considered subspecies of *A. kunkeei* (e.g., *Apilactobacillus nanyangensis, Apilactobacillus waqari, Apilactobacillus sp.* F1*, Lactobacillus sp. M0345* strain) ^26^. A few taxa, such as *Apilactobacillus zhanggiuensis*, showed ANI values of about 90-92% and should be considered different species within the genus *Apilactobacillus* (<u>Table S2</u>). We conclude that none of the 48 smORFs have known homologs outside the genus *Apilactobacillus*.

We attempted expression of 15 of these smORFs from *A. kunkeei* strain A0901 in an *E. coli* vector and were able to purify five of the encoded proteins. For simplicity, we refer to these open reading frames as well as their expressed proteins as smORFs: smORF5 (A0901_04910), smORF7 (A0901_04830), smORF8 (A0901_04820), smORF9 (A0901_04570), and smORF12 (A0901_01190) (<u>Table S1</u>). A certain smORF name, e.g., smORF5, would then represent the orthologs present in strain A0901 and the other strains in which a homolog was found. Three of the smORFs (smORF5, smORF7, smORF8) are located inside a putative phage region and code for very short proteins of 49 to 56 amino acids (Figure S1c). smORF9 is located near to the *cas* gene cassette upstream of the phage region and codes for a protein of 153 amino acids, while smORF12 is located elsewhere and potentially codes for a 66-residue protein (Figure S1c). In addition, we included a short, truncated gene of known origin that encodes a protein of similar size to the smORFs and is a fragment of an IS-element, smORF_IS (A0901_13330). Thus, six proteins were selected in total for more detailed bioinformatics analyses and biophysical experiments.

To investigate whether the six smORFs are expressed under laboratory settings, we collected two proteomics datasets from *A. kunkeei* strain A0901. One dataset was obtained following sampling of bacterial cells during both exponential and stationary growth phases (below referred to as the “log-vs-stat” dataset). The other dataset was obtained following growth under stressful conditions, which were induced by different concentrations of ciprofloxacin that causes replication stalling (below referred to as the “CPX” dataset). Using LC-MS/MS based proteomics analyses, we experimentally identified 712 proteins in the log-vs-stat dataset and 864 proteins in the CPX-dataset. Inspection of the data for the 48 smORFs with no or few hits to other species showed that 11 proteins were identified in at least one dataset (<u>Table S3</u>). These included smORF7 and smORF8, which were identified in both datasets as well as smORF5 and smORF9, which were identified in the CPX-dataset (<u>Table</u> <u>S3</u>). No protein could be detected for smORF12 in either dataset, but it was nonetheless selected for further studies because of its high conservation and apparent non-phage origin. Since the sequence of the smORF_IS is present in multiple copies in the genome its expression pattern cannot be assessed, because the peptides obtained in the LC-MS/MS analyses does not distinguish matches to smORF_IS from matches with all other IS-elements.

### Phyletic distribution profiles of the smORFs

We first inspected the BLAST hits to species outside *A. kunkeei* for the five smORFs of unknown function and origin. The identified smORFs in the phage region (smORF5, smORF7 and smORF8) had no homologs in any other species with sequence identity values above 50%. However, both smORF9 and smORF12 have homologs in *A. zhangqiuensis* with sequence identity values in the range of 74% to 95% (E-values of 2.8e-39 for smORF12 (A0901_01190) and 5.56e-79 for smORF9 (A0901_04570) (see, Table S1). Thus, the three smORFs within the phage region seem to be restricted to *A. kunkeei*, while the other two have homologs in at least one other species within the genus *Apilactobacillus*.

Next, we examined the occurrence of the five smORFs within the *A. kunkeei* population. To this end, we determined the phyletic distribution pattern of the smORFs in 104 *A. kunkeei* isolates for which closed genome sequences are available. To facilitate the downstream, comparative analyses, we selected a set of 34 isolates that represent the phylogenetic diversity of the 104 *A. kunkeei* strains as described previously ^25^ and mapped the phyletic distribution patterns of the smORFs onto the phylogeny (**Figure 1**). However, strain H3B1-11M which was previously used as the reference for a cluster of 12 isolates did not contain any smORFs and was substituted for isolate H4B5-02X from the same cluster, which contained both smORF5 and smORF7. We also included H3B2-03J from this cluster because it contained the entire phage region, including smORF8. Likewise, H3B2-06M, which was used as the reference strain for a cluster of seven isolates, was substituted for strain H4B2-10M, which was the only isolate in that cluster that contained smORF7 and smORF8 (**Figure 1**).

**Figure 1.**
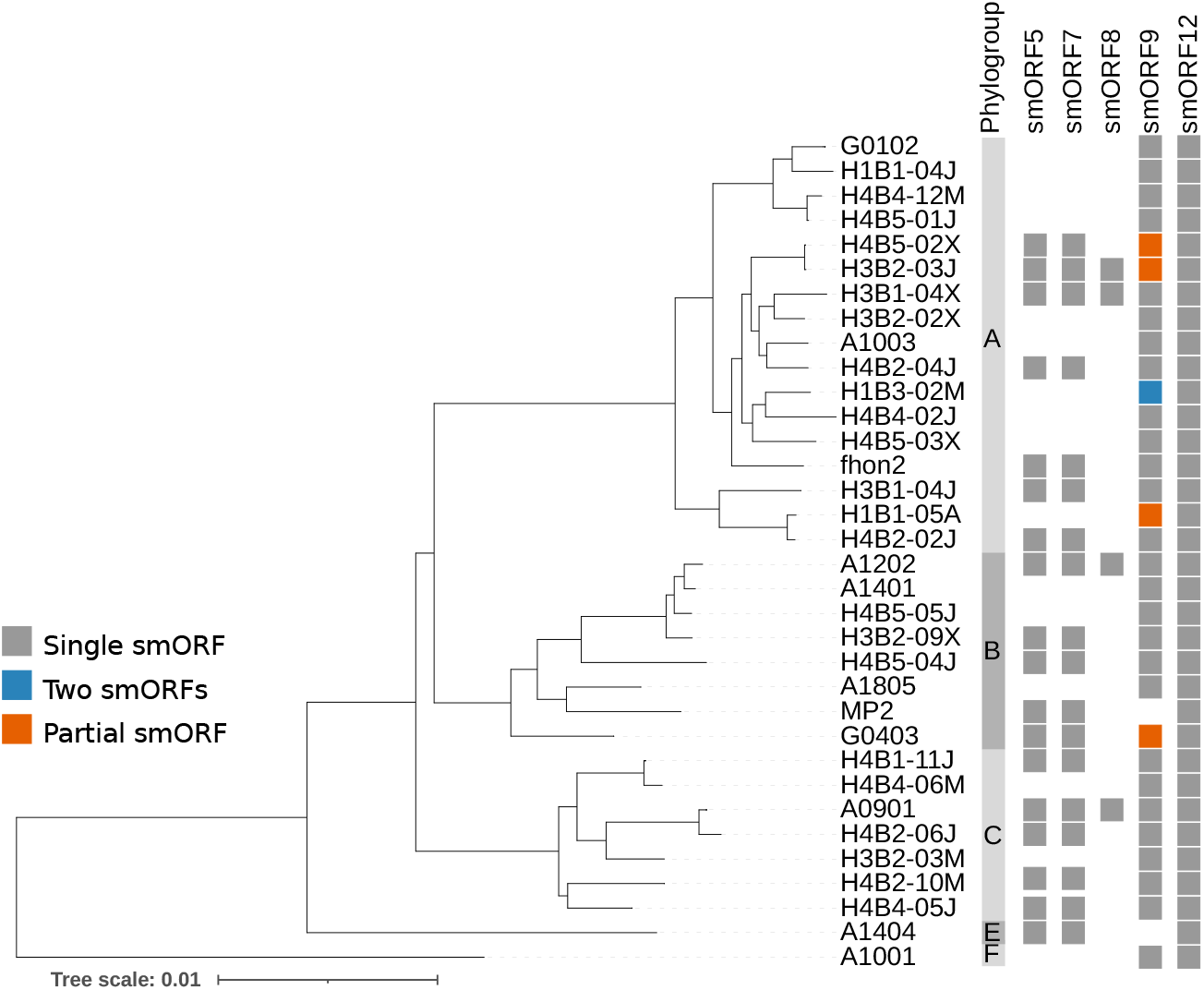
Phyletic distribution patterns of smORFs. The presence and absence pattern of short genes of unknown function among 34 representative *A. kunkeei* isolates. Filled boxes represent the presence of a smORF homolog whereas an empty box indicates that no sequence homolog could be detected in that particular strain. Phylogroup classifications of strains is indicated (Dyrhage et al. 2022).

The analyses showed that two genes were highly conserved within the population; smORF12 was present in all strains and smORF9 in all strains except *A. kunkeei* strains MP2 and A1404 (<u>Figure S2</u>). The other three genes, smORF5, smORF7 and smORF8, were located in close vicinity to each other in the genome (**Figure 2**; Figure S1c), and were predicted to be located within or near a phage region that also contained many other hypothetical genes with no or few hits outside the genus *Apilactobacillus* ^25^. Although we refer to all of these genes as ‘phage genes’, it has not been experimentally verified whether they really code for phage proteins. The phage genes were only present in a subset of strains; smORF5 and smORF7 were present in 40 isolates, of which 11 isolates also contained smORF8 (<u>Figure S2</u>). The strains containing the phage smORFs were phylogenetically disperse suggesting multiple independent phage integration/excision events. This hypothesis was further supported by the finding that clades consisting of *A. kunkeei* isolates with otherwise identical genomes differed with regard to the occurrence of these three phage genes (<u>Figure S2</u>).

**Figure 2.**
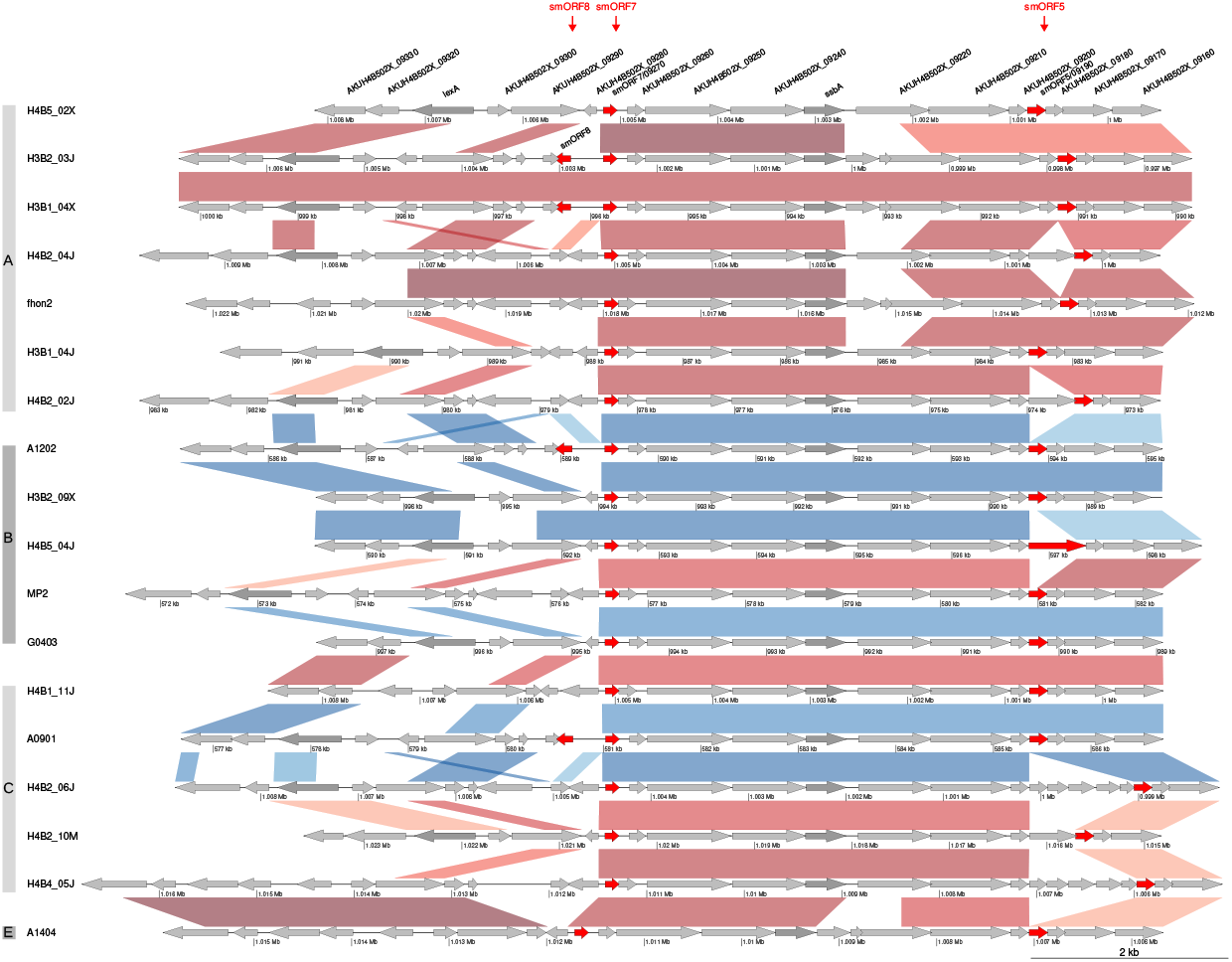
Gene order structures of smORFs located in a phage region. The figure shows a comparison of gene order structures for smORF5, smORF7 and smORF8 in a set of representative *A. kunkeei* strains. The smORFs are highlighted with red arrows, while genes marked in grey and dark grey represent hypothetical and functionally annotated protein-coding genes, respectively. The connecting vertical lines between any two strains show genes and segments with high levels of sequence similarity. The red and blue colors of these lines show segments located at the same or different chromosomal sites, respectively. Phylogroup classifications of strains is indicated (Dyrhage et al. 2022).

The phage segment (containing smORF5, smORF7 and smORF8) has been integrated at either of two chromosomal sites that flank the terminus of replication (ca 500 kb and 900 kb respectively) (**Figure 2**) ^25^. Isolates from Åland contained the phage region at the 500 kb site and this insertion site was also observed in a single isolate obtained from a single bee in hive 4 from Helsingborg (H4B5_04J), whereas all other phage-containing isolates from Helsingborg and Gotland contained the phage insertion at the 900 kb location. The 11 isolates that contained smORF8 belonged to four lineages, two of which contained strains from Helsingborg, while the other two strains were sampled from bees at Åland (**Figure 2** and <u>Figure S2</u>) ^25^. Interestingly, smORF9 was located near to the phage region and the CRISPR-cas segment and was present in all but two genomes (Figure S1c). A single genome contained two copies of this gene (Figure S2c). Finally, smORF12 was located at the same chromosomal site near the origin of replication in all genomes, and as such it is the most broadly distributed smORF among those analyzed here.

### Protein sequence conservation and gene order structures

To learn more about the selective pressures acting on the *A. kunkeei* smORFs, we examined the extent of protein sequence conservation. Overall, the high amino acid conservation of all five smORFs, as detailed below, suggests they are under selection for a function. For example, the putative phage protein smORF5 is highly conserved in sequence among all isolates (<u>Figures S2, S3a</u>). Only five amino acid replacements (in 63 residues) were noted in two reference strains from phylogroup C, one of which represents a cluster of four isolates. A larger change was observed in the strain H4B5_4J, but is not discussed further here (<u>Figure</u> <u>S3a</u>).

The second phage protein, smORF7 (63 amino acid residues), is also highly conserved in amino acid sequence across all isolates, irrespectively of the insertion site of the phage (<u>Figure S3b</u>). smORF7 is the first gene in a long stretch of phage genes in the same transcriptional direction (including the *ssb* gene for the single strand binding protein), and as such likely to correspond to the first gene in a phage operon (**Figure 2**). The alignment reveals only a few single nucleotide polymorphisms (SNPs) in the 5’ and 3’ ends of the smORF7 gene, respectively (<u>Figure S3b</u>). The SNPs do not reflect the phylogenetic relationships of the strains, but indicates intragenomic recombination events and/or integration with a few different phage variants.

The third phage protein, smORF8, is 56 amino acids long and perfectly conserved in sequence among all the eleven isolates in the three different lineages (**Figure 1**, <u>Figure S2,</u> <u>Figure S3c</u>). smORF8 is located immediately next to smORF7, but in the opposite direction, and they are separated by a noncoding region of about 1 kb (**Figure 2**). The eleven isolates containing smORF8 also contains a unique SNP in the N-terminal end of smORF7 not found in any of the other isolates, but differ with regard to a SNP at the C-terminal end (<u>Figures</u> <u>S3b, S3c</u>). This suggests that the recombination or integration events span across both smORF7 and smORF8, with a break-point within the smORF7 gene. In some strains, an ORF of length 192 bp is predicted to be a homolog present in the intergenic region of smORF7-smORF8 (<u>Figure S3d</u>). The gene for the LexA protein is located further away and the segment between smORF7 and the *lexA* gene is hypervariable and represented by more than five different gene order structures. Isolates with a similar gene order structure in this region belong to several different phylogenetic clades, which argue against vertical inheritance and suggest horizontal transmission and exchange.

Multiple sequence alignment of smORF9 (153 residues) homologs revealed the presence of two different sequence variants that differ at multiple sites (<u>Figure S4b</u>). Gene variant I, which is present in *A. kunkeei* strain A0901, is additionally found in most strains of phylogroup C as well as those sampled from Åland in phylogroup F, whilst gene variant II is most common in isolates from phylogroups A and B. The start codon is AUG for methionine in the gene variant I sequences, but CUG for leucine in the gene variant II sequences, although these sequences contain a downstream AUG codon that may alternatively function as the start site. A few isolates in gene variant II contained a short deletion corresponding to six amino acids *(i.e.*, eighteen nucleotides), as well as multiple SNPs and a single nucleotide deletion that disrupts the open reading frame. This raises the possibility that the smORF9 sequences in these isolates code for an even shorter protein that is started from a third AUG codon. Two isolates, Fhon2 and H1B1-05A contain a truncated C-terminal region of smORF9 (<u>Figure S4b</u>). However, inspecting the sequence manually showed a conserved sequence both before and after the stop codon, and we can therefore not exclude that the SNP is a sequencing error.

smORF12, which is present in a single copy in all isolates, is highly conserved in sequence with only a few SNPs, although gene order structures vary widely among the isolates (<u>Figure S5a, b</u>). Finally, smORF_IS is a short fragment of a multi-copy IS element. Transposons are known as jumping genes and in *A. kunkeei* isolates their numbers range from 2 to 90 per genome between different clade groups (see Figure 4 and Figure S3 in Dyrhage et al. ^25^). In the A0901 strain, one of the transposons is truncated and split into two smaller fragments (genes) (<u>Figure S6a</u>). Comparing smORF_IS_A0901 to smORF_IS from the other *A. kunkeei* isolates within the similar genomic region showed that an IS-element was present at this location in seven additional isolates. However, only three of them had smaller fragments similar to smORF_IS_A0901 and with almost identical amino acid sequences (<u>Figure S6a, b</u>).

### The new proteins possess secondary structure

To assess the secondary structure of the proteins, we performed far-UV circular dichroism (CD) experiments 25°C (**Figure 3a**). smORF5, smORF7, smORF12 and smORF_IS displayed a typical alpha helical signature with two minima around 210 and 222 nm, whilst smORF8 and smORF9 displayed CD spectra consistent with a mixed alpha helix/beta sheet structure. One of the proteins, smORF12, appeared as partially unfolded as judged by the CD spectrum. CD spectra at 90°C were consistent with a lower degree of secondary structure as compared to 25°C, suggesting that all six proteins contained secondary structure distinct from a random coil under physiological temperature. SmORF9, which is the largest of the proteins, displayed the lowest molar ellipticity suggesting it may contain more regions lacking secondary structure as compared to the other five proteins.

**Figure 3.**
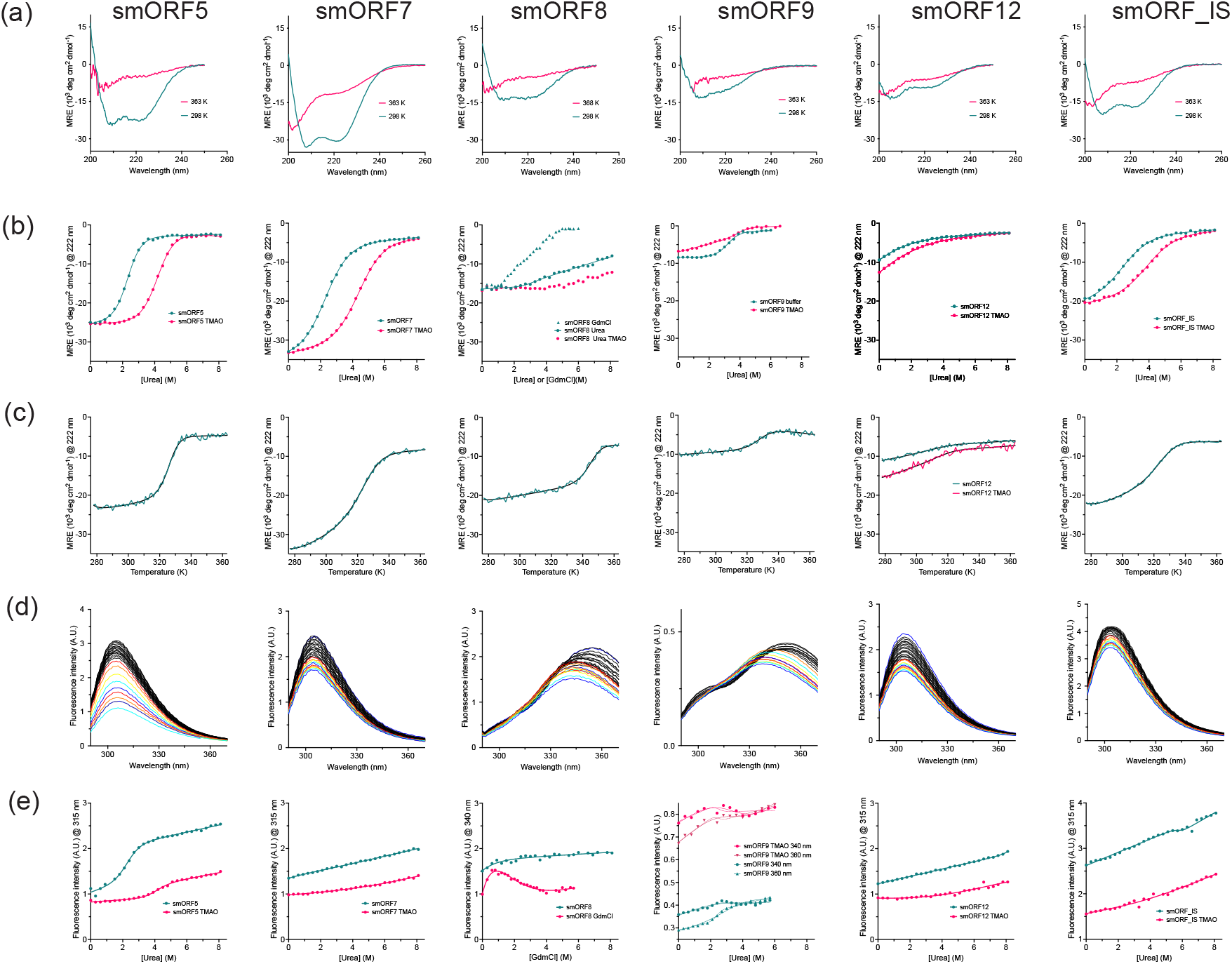
Secondary structure of the new proteins. (a) Far-UV circular dichroism spectra show that the smORF proteins predominantly adopt helical structures. (b) Denaturation experiments using urea or GdmCl monitored by circular dichroism at 222 nm where alpha helical secondary structure gives a strong negative signal. A two-state model was fitted to the experimental data. (c) Heat denaturation experiments monitored by circular dichroism at 222 nm. (d) Fluorescence emission spectra following excitation at 274 nm or 280 nm (smORF8 and smORF9). (e) Denaturation experiments using urea or GdmCl monitored by fluorescence emission. A sigmoidal shape indicates cooperative (un)folding of the protein in panels (b), (c) and (e). Fitted parameters from all experiments are shown in <u>Table S5</u>. Experiments were performed in 50 mM sodium phosphate, pH 7.4.

To further investigate whether the smORF proteins adopt a folded structure we determined their thermodynamic stability with increasing denaturant concentration (urea or guanidinium chloride, GdmCl) (**Figure 3b**) or temperature (**Figure 3c**), respectively. Proteins with well-defined tertiary structures and a hydrophobic core typically unfold in a cooperative, sigmoidal fashion. In such cases, the slope of the transition region (*m*_D-N_ value) is related to the difference in solvent accessible surface area upon unfolding. Thus, a larger protein will generally have a higher *m*_D-N_ value than a smaller protein. Moreover, if the unfolding transition is reversible, the thermodynamic stability of the protein can be estimated by assuming an apparent two-state scenario, were the native folded state and the denatured unfolded state are the only dominating molecular species under all conditions (temperature or denaturant concentration). Urea denaturation experiments, performed both in the presence and absence of the stabilizing molecule trimethylamine *N*-oxide (TMAO) and monitored by far-UV CD at 222 nm, showed clear cooperative unfolding transitions for five of the six proteins tested, corroborating the presence of secondary structure under native conditions (**Figure 3b**). The transition of smORF12 did not contain a native baseline, confirming that the protein is partially unfolded at 25°C. The urea denaturation transition of smORF8 was very broad and incomplete, and therefore, we used the stronger denaturant GdmCl, which resulted in a non-cooperative broad unfolding transition. This behavior is consistent with a multi-step unfolding mechanism involving intermediates present at equilibrium. While smORF9 displayed a single cooperative transition in buffer without TMAO, addition of this stabilizing agent resulted in a broad transition suggesting that a more complex (un)folding process was induced by TMAO. Temperature denaturation experiments recapitulated the denaturant experiments, further demonstrating the presence of secondary structure in the native state (**Figure 3c**). Here, smORF8 displayed an apparent cooperative transition with a relatively high thermal midpoint (72°C) for such a small protein domain.

Next, we performed urea denaturation experiments and monitored fluorescence, which probes gross changes in tertiary structure from the changes in the environment of Trp and Tyr residues upon denaturation. Typically, a change from hydrophobic to solvent exposed environment results in a blue shift of the maximum emission wavelength. However, only smORF8 and smORF9 displayed a clear shift in emission maximum, suggesting that the aromatic residues in the other proteins do not experience an increase in solvent exposure upon denaturation and, thus, are likely not part of a hydrophobic core. Consistent with this result, smORF7, smORF12 and smORF_IS did not display any visible transition when the urea denaturation was monitored at 315 nm (the emission wavelength of Tyr residues). This result could either be interpreted as a lack of globular structure in the native state, or that the Tyr residues are solvent exposed in the native state (as is the case for smORF7, see below). smORF5 displayed a cooperative transition, whereas smORF8 again showed a non-cooperative behavior upon addition of GdmCl consistent with a multi-step unfolding (emission at 340 nm due to presence of one Trp). smORF9 displayed a cooperative transition and, like for the CD data, addition of TMAO complicated the urea dependences with a switch from increase to decrease of the fluorescence upon unfolding at high (360 nm) but not low (340 nm) emission wavelength. The presence of two Trp residues in the sequence of smORF9 underlies the increase or decrease of fluorescence at different emission wavelengths.

### Thermodynamic stabilities of the new proteins are in the typical range of small domains

Naturally occurring protein domains are usually marginally stable with free energies of folding (Δ*G*_D-N_) in the range 2-5 kcal mol^-1^. In order to assess the thermodynamic stability of the *A. kunkeii* smORFs, the urea and temperature denaturation data were analyzed according to a two-state (un)folding scenario where the native and denatured states are assumed to be in equilibrium. From denaturant-induced unfolding data, Δ*G*_D-N_ is calculated from two parameters obtained from fitting a two-state mechanism: the midpoint of denaturation ([Urea]_50%_) and the *m*_D-N_ value, which is related to the change in solvent accessible surface area upon unfolding. In temperature denaturation experiments, Δ*G*_D-N_ is calculated from three fitted parameters: the thermal midpoint (*T*_m_), the change in enthalpy of unfolding (Δ*H*_D-N_) and the change in heat capacity (Δ*C*_p_) (<u>Table S5</u>). While midpoints can usually be determined accurately, the other parameters may have large errors for small proteins where the transitions are broad and baselines not well defined. (In particular, Δ*C*_p_ is very uncertain, but Δ*G*_D-N_ is not very sensitive to errors in this parameter.) Nevertheless, data from the different experiments were overall consistent and show that the smORF proteins display thermodynamic stabilities that are in the same range as typical protein domains of a similar size (<u>Table S5</u>). Δ*G*_D-N_ values derived from CD-monitored urea and temperature-induced denaturation agreed fairly well for smORF5 (2.5-3.3 kcal mol^-1^), smORF7 (1.4-1.9 kcal mol^-1^), smORF9 (4.3-4.7 kcal mol^-1^) and smORF_IS (1.4-1.7 kcal mol^-1^). smORF8 displayed non-cooperative transitions in denaturant-induced experiments but a clear cooperative transition in temperature denaturation (Δ*G*_D-N_ = 3.2±1.6 kcal mol^-1^). smORF9 displayed a transition towards non-cooperative unfolding in presence of TMAO, as monitored both by CD and fluorescence. The thermodynamic parameters (*m*_D-N_ and [Urea]_50%_) from fluorescence-monitored denaturation were particularly non-consistent with and without TMAO. Association of monomers into dimers or higher order quaternary structure may underlie the observed non-cooperativity, both for smORF8 and smORF9. smORF12 was the least stable of the proteins and populated the native state to only around 50% at 25°C (Δ*G*_D-N_ = −0.6 to 1.2 kcal mol^-1^).

### The new proteins display simple tertiary structure

To obtain more detailed structural information we expressed and purified the six proteins as single (^15^N) or double (^13^C/^15^N) labeled samples and subjected them to nuclear magnetic resonance (NMR) spectroscopy experiments. ^1^H-^15^N heteronuclear quantum coherence (HSQC) spectra (**Figure 4**) were well-dispersed and allowing for high resolution NMR spectroscopy analysis. From the NMR experiments, including dihedral angles, ^3^*J* couplings constants and distance information obtained from nuclear Overhauser effects (NOEs), we were able to perform a complete backbone and sidechain NMR assignments and determine the 3D structures of smORF5, smORF7 and smORF_IS at 25°C in 50 mM sodium phosphate pH 6.5 (**Figure 4 and 5)**. The conformations of smORF8 and smORF12 were heterogeneous under these conditions, *i.e.*, the proteins populate more than one conformation, which is consistent with thermodynamic data. smORF9 was poorly expressed in minimal medium and we could not get sufficient amounts for 3D NMR experiments. Nevertheless, our NMR experiments revealed that the smORFs mainly contain helical structures in agreement with the CD data.

**Figure 4.**
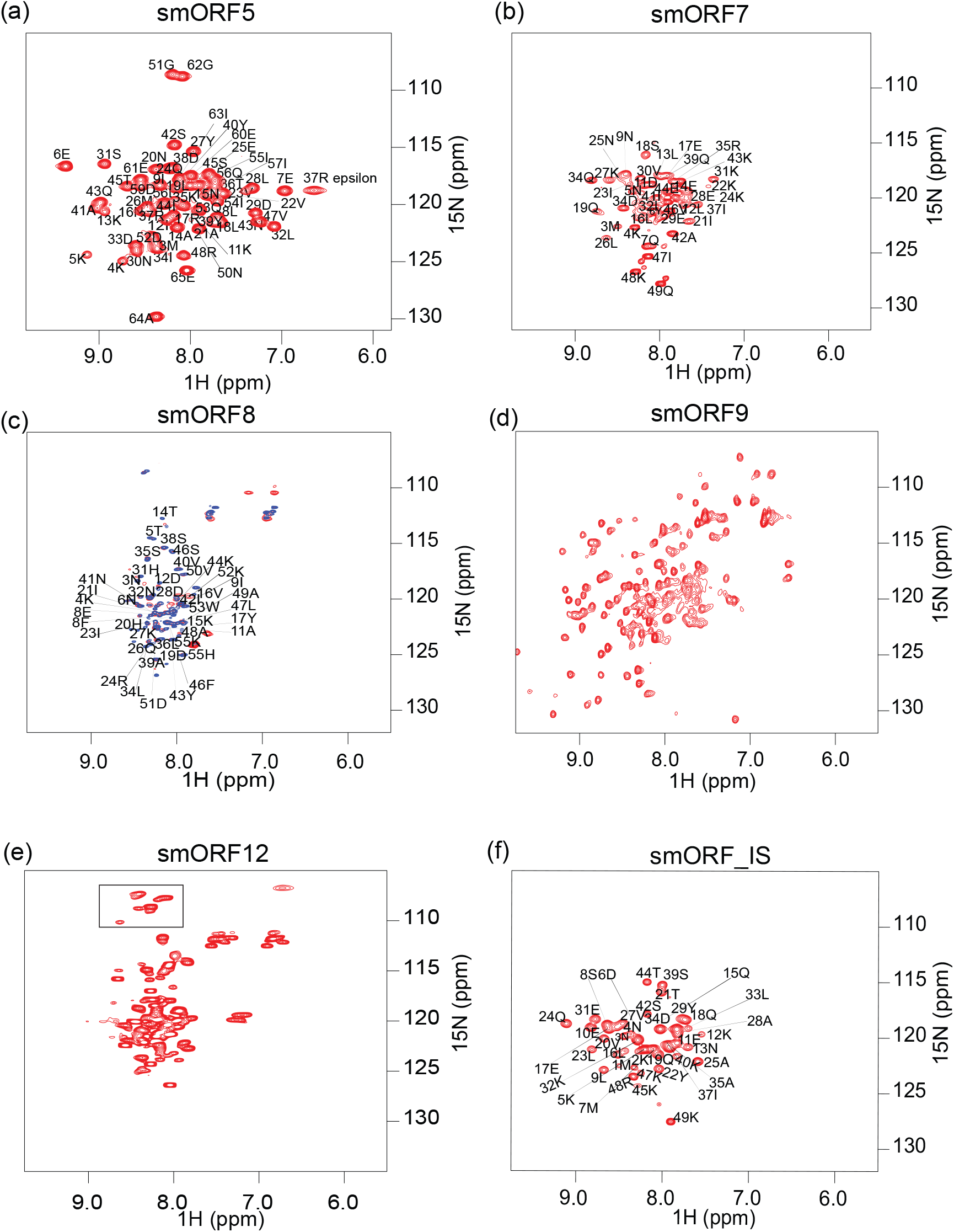
^1^H-^15^N HSQC spectra and backbone assignments of the new proteins. For spectra with amino acid assignment, the side-chain amide resonances of Asn and Gln are omitted for clarity (a) Complete assignment of all ^1^H-^15^N pairs for smORF5. (b) Complete assignment of all ^1^H-^15^N pairs except for proline residues. (c) The assignment of smORF8 was performed for the acid-denatured state at pH 4.0 (blue). The spectrum of native smORF8 at pH 6.0 (red) did not show resonances for all amino acids. However, most of the resonances at pH 4.0 were also visible at pH 6.0 indicating that a state, similar to the acid-denatured one, is present under physiological conditions. (d) The ^1^H-^15^N HSQC spectrum of smORF9 shows that it is well folded and has beta strands because of the spread of the resonances. (e) The ^1^H-^15^N spectrum of smORF12 indicates that it displays structural heterogeneity. 50% of the amino acids were visible and some contain multiple resonances. For example, the resonances above 8 ppm (^1^H) or below 110 ppm (^15^N) (denoted by the rectangle) correspond to the three HN-N pairs of three glycines present in the protein. In a single conformational state only three resonances are supposed to be visible but here six are visible. (f) Complete ^1^H-^15^N assignment of all amino acids in smORF_IS except prolines. The spectrum shows that most peaks exist only in a single state, suggesting one main conformation of smORF_IS.

**Figure 5.**
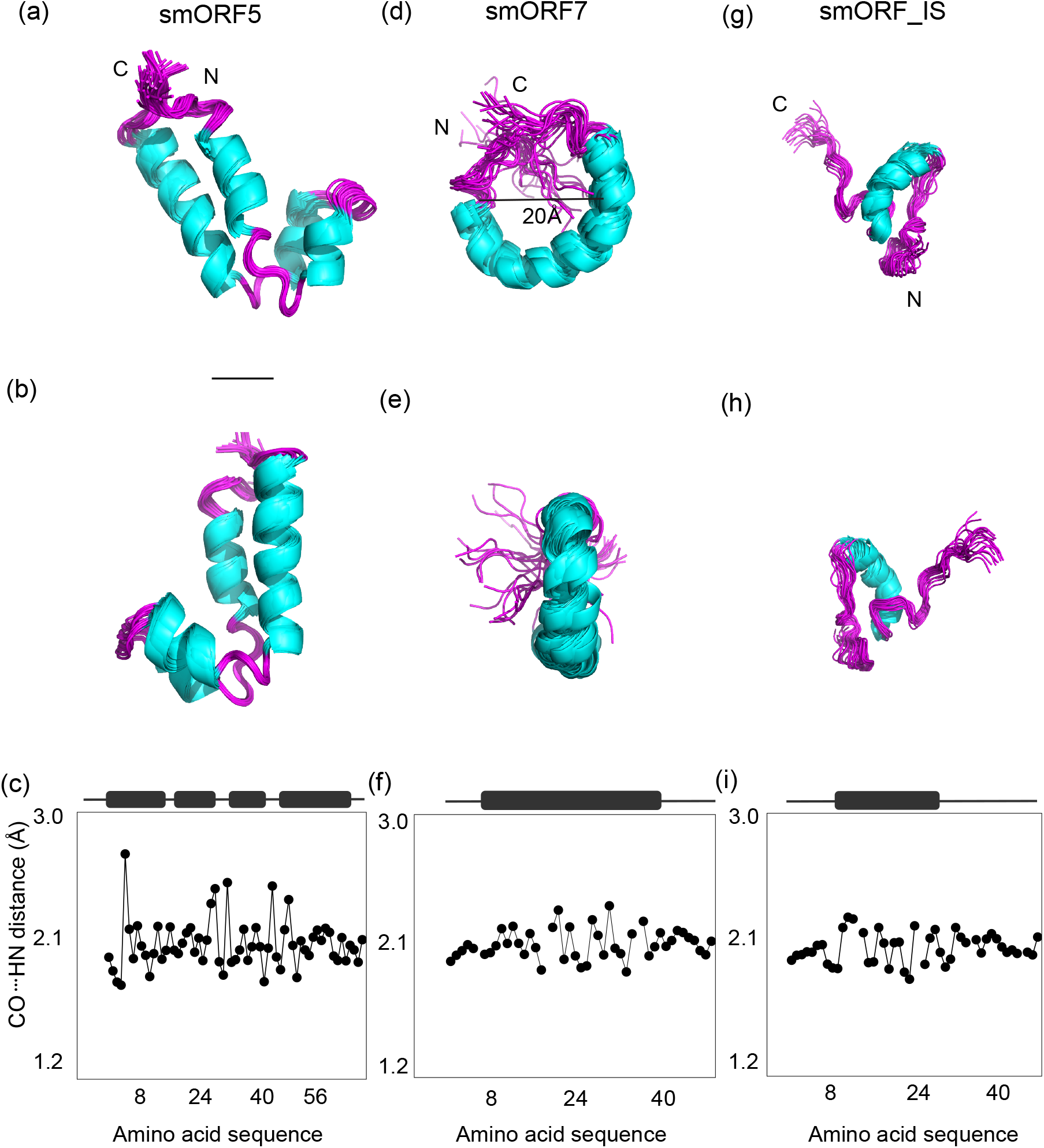
Three dimensional structures of three smORFs and hydrogen bond lengths in their helices. Helices are shown as cyan and loops in purple. (a) and (b) Cartoon representation of smORF5 shows a four-helix bundle including a helix-loop-helix motif, with the N and C termini close to each other. (c) amide (HN) - carbonyl oxygen (CO) bond length determined from ^1^H-^15^N chemical shifts of smORF5. Firstly, the bond lengths follow a repetitive pattern of four amino acids, a feature typical for helices. Secondly, the bond lengths are not uniform along the sequence indicating that the helices are not straight but bent. (d) and (e) The NMR data of smORF7 are consistent with a disc-like fold. Firstly, the data indicate that the protein is well folded, but we observed only a few long-range NOEs. Secondly, the secondary structure propensity predicted from the chemical shifts indicates a single helix. Thirdly, the CO-HN bond lengths determined from HN-H spectra (f) suggest that the helix is not straight. Together these observations point to a circular structure, similar to that of membrane associated proteins of so-called nano discs, and consistent with short-range but not long-range NOEs. Furthermore, T1/T2 relaxation data suggest that smORF7 is a dimer. Finally, the N- and C-termini are not well defined. This could be partly due to the few structural restraints, but also to a high flexibility in this region. The diameter of the disc is measured to 20 Å. (g) and (h) Cartoon representation of smORF_IS. This protein adopts a single short helix, which makes long-range interactions with parts of the unstructured N- and C-termini. (i) The CO-HN bond pattern for smORF_IS indicates that the folded helix is not straight.

To probe the nature of the helices, we determined the hydrogen bond lengths between the amide proton (HN) and carbonyl oxygen (CO) atom for smORF5, smORF7 and smORF_IS using chemical shifts of HN from the ^1^H-^15^N HSQC experiment (**Figure 4**). In alpha helices a hydrogen bond is formed between HN of amino acid *i* with the carbonyl oxygen of an amino acid at the *i*+4 position. These bond lengths range between 2.8-3.2 Å. Hydrophobic pairs accumulate in the interior of the protein resulting in shorter hydrogen bond lengths while hydrophilic pairs on the exterior have longer hydrogen bond lengths. This arrangement will result in a 3-4 periodicity repeat and a curvature of the helix with the longer hydrogen bonds outside and the shorter ones inside. On the other hand, a mixture of hydrophobic-hydrophilic pairs will result in an average bond length similar across the chain. We found that the helices in smORF5, smORF7 and smORF_IS displayed the 3-4 repeat periodicity and the bond lengths alternating from short to long and longer stretches of shorter hydrogen bond lengths indicating bent structures ^27^. smORF5 folds into two kinked parallel helices with the N and the C termini very close to each other. We observed several long-range NOEs (<u>Figure S7 and S8</u>) making the structure of smORF5 the most well defined of the three we determined. smORF7 appears disc-shaped with a diameter of approximately 20 Å. Only a few long-range NOEs were observed for smORF7 (<u>Figure S9 and S10</u>), which is typical for a circular helical structure ^28^. On the other hand, smORF_IS folds into a single helix with intrinsically disordered N and C termini. Interestingly, smORF_IS contains several long-range NOEs between the ordered region and both the disordered N-terminus and C terminus (<u>Figure</u> <u>S11 and S12</u>). More specifically, the ordered helix stretches between residues Glu17-Lys32. We observed NOEs between Tyr29 and Gln38, Tyr29 and Glu31, Asn13 and Ala35, Asn3 and Lys32, Lys2 and Lys32, Leu30 and Asp34, Tyr29 and Leu33, and Leu9 and Lys32, suggesting that the disordered C and N-termini make transient interactions with the ordered central helix.

NMR relaxation parameters harbor several indicators of the behavior of the protein such as flexibility, shape and size. For example, NMR relaxation parameters such as longitudinal and transverse times, T1 and T2, respectively, give dynamic information in the ps to ms timescale but can also be used to estimate the molecular size. The ratio of the relaxation times (T1/T2) gives an estimate of the total rotational correlation time (τ_C_), which is directly proportional to molecular size and can give a good estimate of molecular mass (**Figure 6**). The theoretical τ_C_ values calculated from the molecular masses approximated to 4.9 ns (smORF5, 7.4 kDa), 3.9 ns (smORF7, 5.9 kDa) and 6.6 ns (smORF_IS, 9.9 kDa). The experimental τ_C_ values were 10.9 ns (smORF5), 10.4 ns (smORF7) and 6.3 ns (smORF_IS). Thus, the T1/T2 relaxation data are most consistent with dimeric quaternary structures of smORF5 and smORF7, and a monomeric smORF_IS.

**Figure 6.**
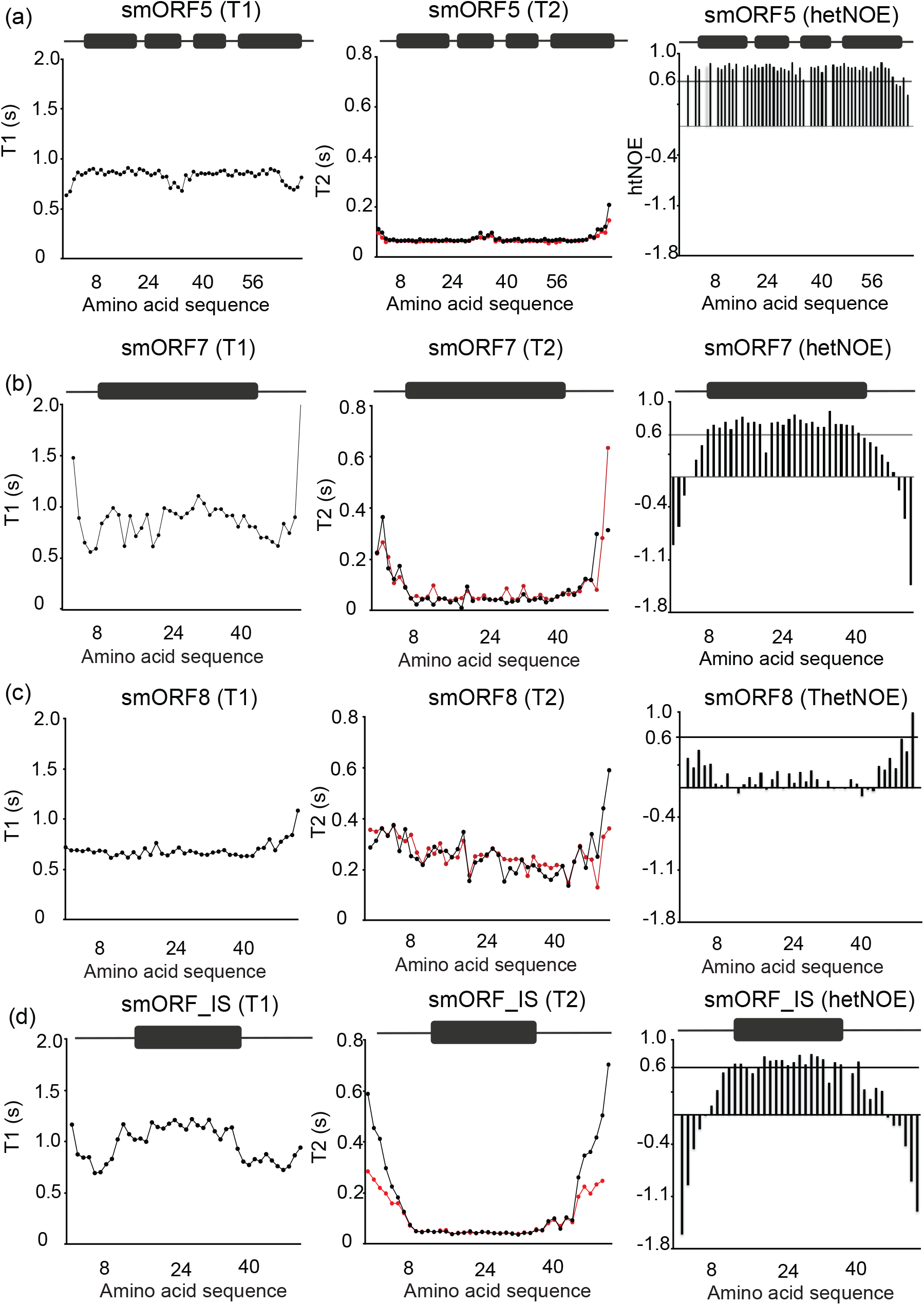
NMR relaxation parameters for the new proteins. The relaxation times and heteronculear NOEs for (a) smORF5, (b) smORF7, (c) smORF8 and (d) smORF_IS. T1 represents the longitudinal relaxation times while T2 represents the transverse relaxation times. hetNOE is the heteronuclear NOE between the amide (HN) hydrogen and the directly bonded nitrogen atom. The combined behavior of these relaxation parameters gives information about ps-ms timescale motion of the protein. The ratio of T1/T2 is unitless, but can be used as an approximation of the total rotational correlation time τ_C_ in nanoseconds, which in turn is related to the molecular mass of the protein.

NMR data collected for smORF8 at pH 6.0 indicated that the protein populated at least two states, where one might be the denatured state under physiological conditions. However, in light of the non-cooperative unfolding transitions, as monitored by CD (**Figure 3b**) and fluorescence (**Figure 3e**), the observed heterogeneity may result from two or more folded states in equilibrium. CD experiments showed that smORF8 is unfolded at pH 4.0. Consistent with this, NMR data at pH 4.0 suggested a single state representing the acid-denatured state of the protein. For smORF12, the observed heterogeneity in the NMR experiments is most likely due to its marginal stability, with around 50 % denatured state under native conditions, as suggested by the CD-monitored denaturation data, which show the second part of an apparently cooperative unfolding transition (**Figure 3b**).

## DISCUSSION

Convergent evolution of phenotypic traits is ubiquitous in nature ^29^. On the other hand, molecular evolution is generally viewed as a result of divergence as shown for the ancient nucleoside-binding Rossmann fold ^30^. In the latter view, contingency is important since new function arises from point mutation or recombination of existing genes encoding a particular unique protein state ^31, 32^. Expressed proteins with a new or modified function would then be based on ancestral protein folds and fine-tuned by adaptation according to changes in selection pressure over evolutionary timescales. However, four key findings during the last decades suggest that emergence of entirely new proteins may be more common than previously thought: (*i*) Intrinsically disordered proteins without a well-defined 3D structure play pivotal roles in the cell ^33^ and are suggested to be more prevalent in young genes ^18^. (*ii*) Non-coding regions are frequently transcribed and translated resulting in small proteins with unknown functions ^4^. (*iii*) New proteins from random libraries can provide a selective advantage in experimental *in vivo* systems ^9–11^. (iv) Globular protein domains can emerge from fusion of small structural motifs ^14, 34–36^. Thus, a consensus is developing that new proteins constantly emerge from novel open reading frames in living organisms ^15, 37^. Usually, the genes encoding these new proteins erode and become non-functional with time, but sometimes they prove beneficial for the organism, such that they are retained in the genome by positive selection. From a protein structure-function point-of-view, a fundamental question is what these new proteins look like. Is it possible that protein folds appear *de novo* such that convergent evolution occur on a molecular structure level, as observed for organismal phenotypes? Predictions suggest that new proteins are largely intrinsically disordered ^18^, but there is a paucity of experimental data regarding their structures.

To shed light on this question we investigated in detail six small proteins from *A. kunkeii* strain A0901 and where five appear to represent young and putative new proteins. The genes were randomly sampled from a larger subset of genes that could be expressed in *E. coli* and for which no homologs were identified outside the genus *Apilactobacillus*. Two genes, smORF9 and smORF12, were successfully identified in all but two *A. kunkeei* strains, suggesting that they represent novel, functional genes. The other three genes, smORF5, smORF7 and smORF8, were located within or near a previously identified prophage region of 35-50 kb that displayed a scattered phyletic distribution pattern within the population. Our analyses of these genes provided indications of frequent integration, excision and recombination events within the population. Although it cannot be excluded that putative homologs might exist also in other bacterial genera, but were missed in our survey due to the short gene lengths and rapid sequence divergence, we consider this possibility unlikely since no or few hits outside the genus *Apilactobacillus* were obtained for any of the longer genes in the prophage region. Importantly, different regions of the bacterial genome may have different likelihood to give birth to new genes, depending on differences in mutation rates, recombination frequencies, proximity to transposons and promoter sequences, etc. For all of these reasons, prophage regions may serve as bacterial testbeds for the evolution of novel genes. Thus, it may not be a coincidence that several of the genes in *A. kunkeei* for which no homologs were found were located within or near to phage regions. In the future, large-scale comparative studies of bacterial genomes are needed to determine the role that phages and viruses play for the origin of novel genes.

We find that the new smORF proteins and smORF_IS from *A. kunkeii* populate simple, mainly alpha-helical folds, and that they are present as monomers or dimers. Five of the proteins are marginally stable, similarly to the numerous small protein domains found across organismal proteomes. (Note that the temperature of the natural habitat of *A. kunkeii*, a bee gut, may greatly vary over the course of a day and over seasons, and it is not known if and how such variation in temperature affects protein evolution and thermodynamic stability.) Two of the proteins, smORF5 and smORF7, are likely present as dimers based on NMR relaxation data. smORF12 is partially unfolded under physiological temperatures, but displays a cooperative transition to a fully unfolded state. smORF7 displays clear cooperative unfolding indicative of a small globular protein, but it has very dynamic N- and C-termini, which nonetheless contribute to the tertiary structure. smORF8 displays rather peculiar biophysical characteristics including a relatively high thermal stability and structural heterogeneity, unlike globular folds of similar size. smORF9 behaves as a two-state folder with less helical content than the other proteins and addition of TMAO induces folding heterogeneity, perhaps by stabilizing cryptic non-native conformational states. Interestingly, we note that none of the five new smORF proteins or smORF_IS behave as a *bona fide* intrinsically disordered protein, but instead display well-defined secondary and simple tertiary structure. A recent high-throughput study, which used limited proteolysis to assess structure in putative *de novo* human and *Drosophila melanogaster* proteins, found evidence for both disordered and ordered proteins in the data set ^38^. Our small set of six smORFs may be strongly biased, but there could also be a difference between eukaryotic and bacterial proteins since disorder is less prevalent among the latter. We note that small *de novo* proteins selected for function from random libraries are predicted to be helical ^9–11^, but that the yeast *de novo* protein Bsc4 contains β sheets as judged by circular dichroism ^39^.

Small new proteins such as the ones investigated here appear to be prevalent across life ^40^. Moreover, the symmetry of many extant protein folds suggest that fusion of small structural motifs is a common pathway for emergence of larger proteins ^35^. The potential mechanism and underlying selective forces were investigated for a lectin β-propeller protein, which consist of ancestral motifs shorter than 50 amino acid residues. Interestingly, a reconstruction of the evolutionary trajectory suggested that constraints related to folding governed the process ^41^, which raises the possibility of structural convergence in new proteins. The new proteins from *A. kunkeii* are based on helical and possibly mixed alpha/beta secondary structure and they display variation in stability, dynamics and quaternary structure. Intrinsic disorder is less prevalent than might be expected *a priori*. Our data suggest that foldable structural motifs may arise continuously in living cells. Such motifs could act as intermediates on a trajectory towards larger *de novo* globular proteins by duplication and fusion of the genes encoding the motifs. Indeed, evolution of protein domains from accretion of smaller sub domains represents a likely mechanism for invention of new domains during origin of life ^14, 16, 42, 43^. Our smORFs may represent such sub domains in an ongoing emergence of *de novo* protein domains, which incidentally could converge on ancient folds. We find it likely that selection for foldability, as part of functionality, may evolve new proteins into similar structures as ancient protein folds. Thus, our present findings and previous data ^35, 41^ suggest that convergent evolution of protein structure is a realistic possibility.

## METHODS

### BLAST search analysis

All the predicted genes (1466) in A0901 strain were subjected to blastp (ver 2.11.0+) against the NCBI NR-database (ver 2023-01-10) with an E-value 0.001 and run with the default parameters ^25, 44, 45^. Identification of orthologous genes among *A. kunkeei* isolates was obtained based on the tblastn analysis against all *A. kunkeei* genome sequences. Blast hits related to any unannotated sequence were manually inspected and included in the multiple sequence alignment.

The chromosome representation of *A. kunkeei* A0901 strain was generated using Circos ver 0.69-9 ^46^. All the gene synteny plots were generated with the R-package genoPlotR ver 0.8.11^47^. Multiple sequence alignments were performed using ClustalWS ver 2.1 ^48^ and visualized with the tool Jalview ver 2.11.2.6 ^49^.

### Proteomic analysis of *A. kunkeei* A0901 under conditions of replication stalling

We collected two proteomics datasets from *A. kunkeei* strain A0901 to investigate whether the predicted smORF genes are expressed under laboratory conditions. The first dataset was obtained by comparing protein expression during exponential and stationary growth phase (“log-vs-stat”). The second dataset was collected during exponential growth phase under the influence of increasing concentrations of ciprofloxacin (“CPX”).

For the log-vs-stat dataset, *A. kunkeei* A0901 was cultivated in MRS broth (Sigma Aldrich) supplemented with Tween-80 and D-Fructose to final concentrations of 0.1% and 0.5%, respectively (referred to as fMRS). Cells were cultivated in biological triplicates at 35°C, 5% CO2 under static conditions and samples were harvested during exponential (Optical density at 600 nm, OD_600_ ≈ 0.2, 3.5 h) and stationary growth (OD_600_ ≈ 1.6, 6 h). Cells were pelleted by centrifugation (3000 *g*, 10 min, 4°C), the supernatant was discarded and the pellet was washed twice in 50 mM Tris, pH 8.0. For protein extraction, the pellets were re-suspended in 25 mM Tris, pH 8.0, 10 µg/mL lysozyme (Sigma Aldrich), 1x Sigma Fast Protease inhibitors (Sigma Aldrich) to achieve a final cell density of OD_600_ ≈ 10 and incubated at 37°C for 5 h under gentle orbital shaking. Additional lysis was achieved by sonication using a Sonics VCX 130 sonicator (Sonics & Materials Inc., 2 mm tip, 4 x 5 s, 5 s pause, on ice). The cell suspension was cleared by centrifugation (16 000 *g*, 10 min, 4°C) and the supernatant was transferred to 1.5 mL reaction tubes. Total protein concentration was determined by Bradford analysis (Thermo Scientific).

For the CPX-dataset, *A. kunkeei* A0901 was cultivated in fRMS broth in the presence of 0.0, 3.1, 6.3 and 12.5 µg/mL ciprofloxacin (CPX, Sigma Aldrich). Batch-cultivation of biological triplicates per condition was performed under static conditions at 35°C, 5% CO_2_ and samples were harvested by centrifugation (4500 *g*, 10 min, 4°C). Pellets were washed twice in HyClone (Cytiva). For protein extraction, washed pellets were re-suspended in 25 mM Tris, pH 8.0, 15 µg/mL Lysozyme and 1x SigmaFast protease inhibitors and incubated at 37°C under gentle orbital shaking for 2.5 h followed by sonication (VCX 130 sonicator, 4 x 5s, 5s pause, 2 mm tip, on ice). The suspension was cleared by centrifugation (16 000 *g*, 10 min, 4°C) and the supernatant was transferred to 1.5 mL reaction tubes. Protein concentration was determined by using the Bradford assay (Thermo Scientific) with BSA as the standard.

Proteomic analysis of the log-vs-stat and CPX-samples was essentially performed as described previously ^50^. Aliquots corresponding to 20 μg protein were taken out for digestion. The proteins were reduced, alkylated and in-solution digested by trypsin according to a standard operating procedure. Thereafter the samples were purified by Pierce C18 Spin Columns (Thermo Scientific) and dried. Dried peptides were resolved in 60 μL of 0.1% FA and further diluted 4 times (CPX) and 5 times (log-vs-stat) prior to nano-LC-MS/MS. The resulting peptides were separated in reversed-phase on a C18-column, applying a 90 min long gradient, and electrosprayed on-line to a QEx-Orbitrap mass spectrometer (Thermo Finnigan). Tandem mass spectrometry was performed applying HCD. Database searches were performed in the MaxQuant software (version 1.5.1.2) ^51, 52^. Proteins were identified by searching against the annotated genome of *A. kunkeei* A0901 ^16^. Fixed modification was carbamidomethyl (C), and variable modifications were oxidation (M), and deamidation (NQ). A decoy search database, including common contaminants and a reverse database, was used to estimate the identification false discovery rate (FDR). An FDR of 1% was accepted for peptides and protein identification. The criteria for protein identification were set to at least two identified peptides per protein.

### Heterologous protein expression and purification

The DNA sequences of all smORF proteins were synthesized and subcloned into a modified pRSET vector by GenScript (Hong Kong). The construct was tailored to have an N-terminal hexa-histidine-tagged lipoyl fusion protein followed by a thrombin cleavage site and the respective smORF protein. The DNA sequences were confirmed by Sanger sequencing (Eurofins Genomics, Uppsala). The smORF8 (A0901_04820) construct encoded only the His-tag and the protein due to problems with thrombin cleavage and purification. *Escherichia coli* BL21(DE3) pLysS cells (Invitrogen) were transformed with the plasmid and grown in 2×TY medium containing 100 μg/mL ampicillin at 37°C. At an OD_600_ of around 0.6, expression was induced by 1 mM isopropyl-β-thiogalactopyranoside. The cells were then grown overnight at 18°C, spun down in a centrifuge at 4°C and resuspended in 20 mM Tris (pH 8.0), 500 mM NaCl. Cells were disrupted by ultrasonication and cell debris was removed by centrifugation at 20,000 *g* for 60 min followed by filtration (0.2 or 0.45 μm). The general purification strategy was as follows: (i) Nickel (II) affinity chromatography (Ni Sepharose 6 Fast Flow, GE Healthcare) in 25 mM Tris (pH 8.0), 500 mM NaCl and 20 mM Imidazole. Bound proteins were eluted by 300 mM or 500 mM imidazole in 25 mM Tris (pH8.0). (ii) Dialysis into 25 mM Tris (pH 8.0), 150 mM NaCl followed by thrombin (GE Healthcare) digestion to remove the lipoyl domain. (iii) A second nickel (II) affinity chromatography step, with 20 mM imidazole included in the washing buffer and where the lipoyl domain binds and the smORF protein is collected in the unbound fraction. (iv) A final purification step involving either ion exchange, size exclusion or reversed phase chromatography. smORF5 (A0901_04910), smORF8 (A0901_04820) (with His tag), and smORF12 (A0901_01190) were purified using a reversed-phase chromatography column (Vydac C8, Grace Davison Discovery Sciences) as the final purification step. The column was equilibrated with 0.1% trifluoroacetic acid and bound proteins were eluted with a 0-100% gradient of acetonitrile (0.1% trifluoroacetic acid). The thrombin-digested smORF7 (A0901_04830) and smORF_IS (A0901_13330) samples were dialyzed against 25 mM Tris (pH 8.0) and loaded onto a Q column (HiTrap Q Fast Flow, GE Healthcare) equilibrated with the same buffer. The smORF proteins eluted in the unbound fraction before the start of a gradient 0-600 mM NaCl in 25 mM Tris (pH 7.5). Although expected to bind an S column, a cation exchanger, neither smORF7 (A0901_04830) or smORF_IS (A0901_13330) did; Q was then used since it bound more impurities. The thrombin-digested smORF9 (A0901_04570) was dialyzed against 25 mM Tris (pH 7.5), 150 mM NaCl, 4 mM DTT, concentrated using Vivaspin columns (Sartorius) and loaded onto a size exclusion chromatography column (S-100, GE Healthcare). Protein purity was checked by SDS-PAGE and the identity of the purified proteins was confirmed by MALDI-TOF mass spectrometry.

### Circular dichroism and fluorescence spectroscopy

Circular dichroism (CD) and fluorescence spectroscopy experiments were carried out on a JASCO J-1500 spectrophotometer and with a Peltier temperature control system at temperatures indicated in the figures. All the CD experiments were performed using a 1 mm quartz cuvette. Protein concentrations were 6-25 μM and the buffer 50 mM sodium phosphate, pH 7.4, unless otherwise indicated. CD spectra were averages of three to five individual spectra. For both chemical (urea or GdmCl) and thermal unfolding experiments, the CD signal was monitored at 222 nm, and a scan speed of 1 K min^-1^ was used for thermal denaturation. Fluorescence spectroscopy experiments were performed with protein in 50 mM sodium phosphate, pH 7.4 in a 10 mm quartz cuvette at 25°C. Emission spectra were recorded with an excitation at 276 nm for smORF5 (35 μM), smORF7 (50 μM), smORF12 (35 μM) and smORF_IS (35 μM), which lack any Trp residues. An exciation wavelength of 280 nm was used for smORF8 (25 μM) and smORF9 (2.5 μM), which contain one and two Trp residues, respectively. In urea and GdmCl-induced unfolding experiments, the emission at 315 nm (Tyr) or 340 nm (Trp, smORF8 and smORF9) was plotted versus denaturant concentration. The (un)folding experiments were analyzed with GraphPad Prism using the standard equations based on a two-state assumption with only native and denatured state significantly populated at equilibrium ^53^.

### NMR spectroscopy

NMR spectroscopy experiments were performed on a Bruker 600 MHz NeoAdvance HD spectrometer equipped with a cryogenic TCI probe (CRPHe TR-1H and 19F/13C/15N 5mm- EZ). smORF proteins were either single (^15^N) or double (^13^C,^15^N) labeled for NMR experiments, by expression in *E. coli* in M9 minimal medium supplemented with 1 g ^13^C glucose and/or 1 g ^15^N ammonium chloride per liter medium. Purification of the labeled proteins were as described for unlabeled proteins. The protein concentration for assignment and subsequent structure determination ranged from 0.5 mM to 2 mM. For smORF5, smORF7, smORF8 and smORF_IS, the following NMR experiments were recorded for backbone and side-chain assignment on a double-labeled ^13^C-^15^N protein sample: ^1^H-^15^N HSQC, ^1^H-^13^C HSQC, HNCACB, HNCoCACB, HBHACoHN, HNCA, HNCoCA, ^1^H-^1^H ^13^C resolved HCCH-TOCSY. For smORF9 only ^1^H-^15^N HSQC was recorded on a ^15^N labeled sample. While for smORF12 ^1^H-^15^N HSQC was recorded on a ^13^C-^15^N labeled sample. The following were the buffer composition and measuring temperatures for the different protein samples: smORF5, smORF7, smORF9 and smORF_IS were measured in 50 mM sodium phosphate pH 6.5 at 298 K, smORF8 was measured in 50 mM sodium acetate pH 4.0 at 298-315 K and smORF12 in 20 mM sodium phosphate, pH 6.0, at 298 K. Protein samples were supplemented in 10% D_2_O, 0.1 % sodium azide. ^15^N and ^13^C resolved ^1^H-^1^H NOESY (28 (15N or 13C) x 256 (^1^H) x 2048 (^1^H, direct)) were measured with mixing times ranging from 70-120 ms and used for distance estimation during structure determination. ^3^*J*_HNHA_ couplings used for structure calculations were measured with a 3D HNHA type experiment. Phi-Psi dihedral angles were estimated using TALOSN. Structure calculation was done with CYANA 3.98 by simulated annealing in 10,000 steps. A total of 100 conformers were calculated and 20 with the lowest target function were selected for analysis. For relaxation experiments, T1, T2, T1rho and heteronuclear NOE (hetNOE), were estimated using standard Bruker pulse programs using randomized relaxation delays of 7-10 durations. The D1 delay was set to 3-5 seconds. The rotational correlation time τ_C_ was estimated from the ratio of T1/T2. Data were evaluated with Topspin and the Bruker program DynamicCenter 2.8.01. All other experiments were processed with Topspin version 4 series. Assignments were performed in the CcpNmr analysis software.

## Acknowledgements

This work was supported by the Swedish Research Council (2020-04395 to PJ and 2018-4135 to SGEA), and Knut and Alice Wallenberg foundation (2015.0069 to SGEA and PJ; and 2018.044 and 2020.0305 to SGEA). We used the NMR Uppsala infrastructure, which is funded by the Department of Chemistry - BMC and the Disciplinary Domain of Medicine and Pharmacy. The computations were performed on resources provided by the National Academic Infrastructure for Supercomputing in Sweden (NAISS) at UPPMAX under project NAISS 2023/22-319 partially funded by the Swedish Research Council through grant agreement no. 2022-06725.

## Author contributions

Conceptualization: SGEA, PJ; Methodology: CNC; Investigation: WY, BPRK, KD, CS, JDJ, EK, EA, CNC; Data acquisition and analysis: WY, BPRK, KD, CS, JJ, EK, CNC; Funding acquisition: SGEA, PJ; Project administration: PJ; Supervision: SGEA, PJ; Writing - original draft: SGEA, PJ; Writing: review and editing: WY, BPRK, JDJ, CNC, SGEA, PJ

## Supplementary Information

### Supplementary Figures

**Figure S1.**
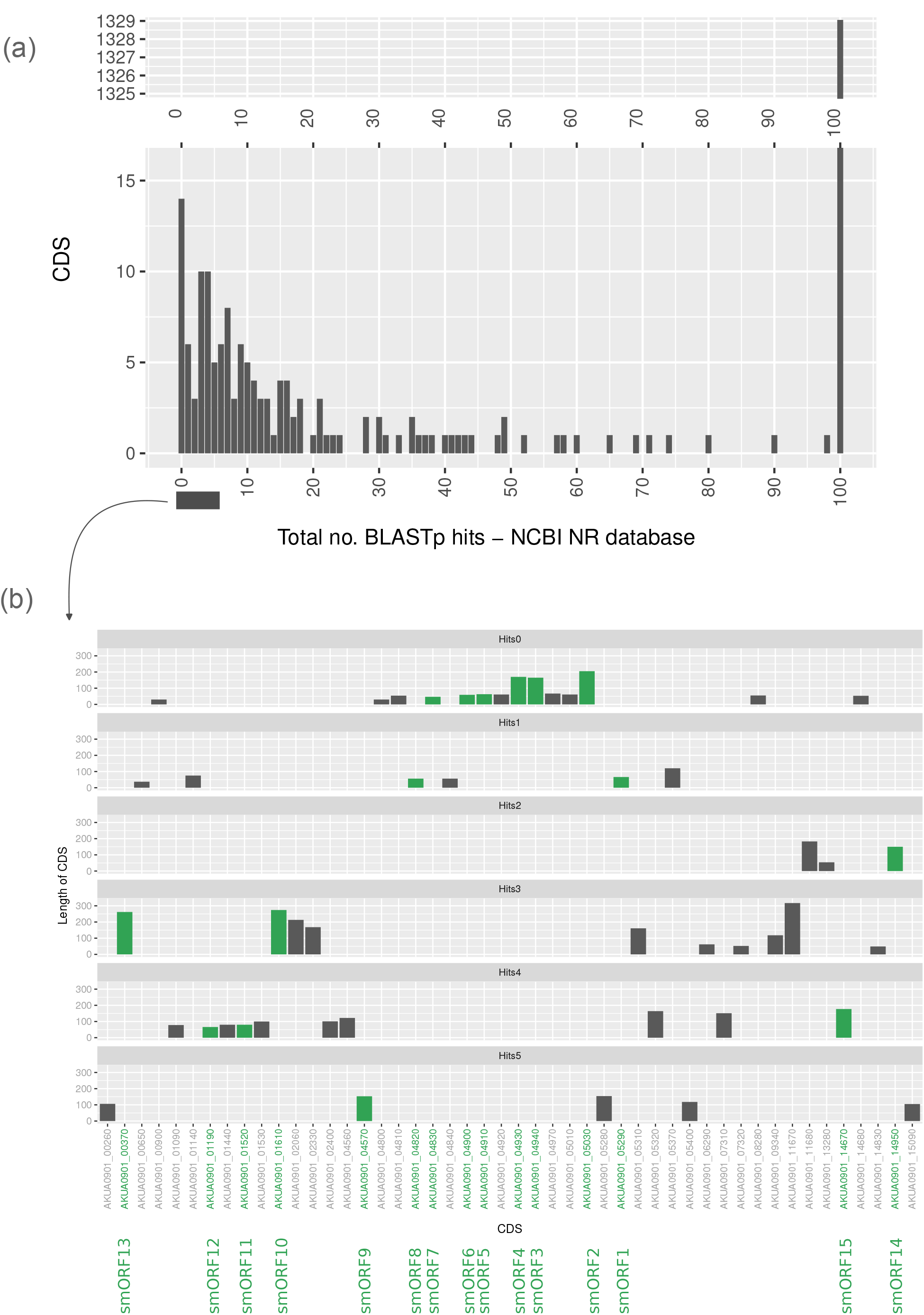
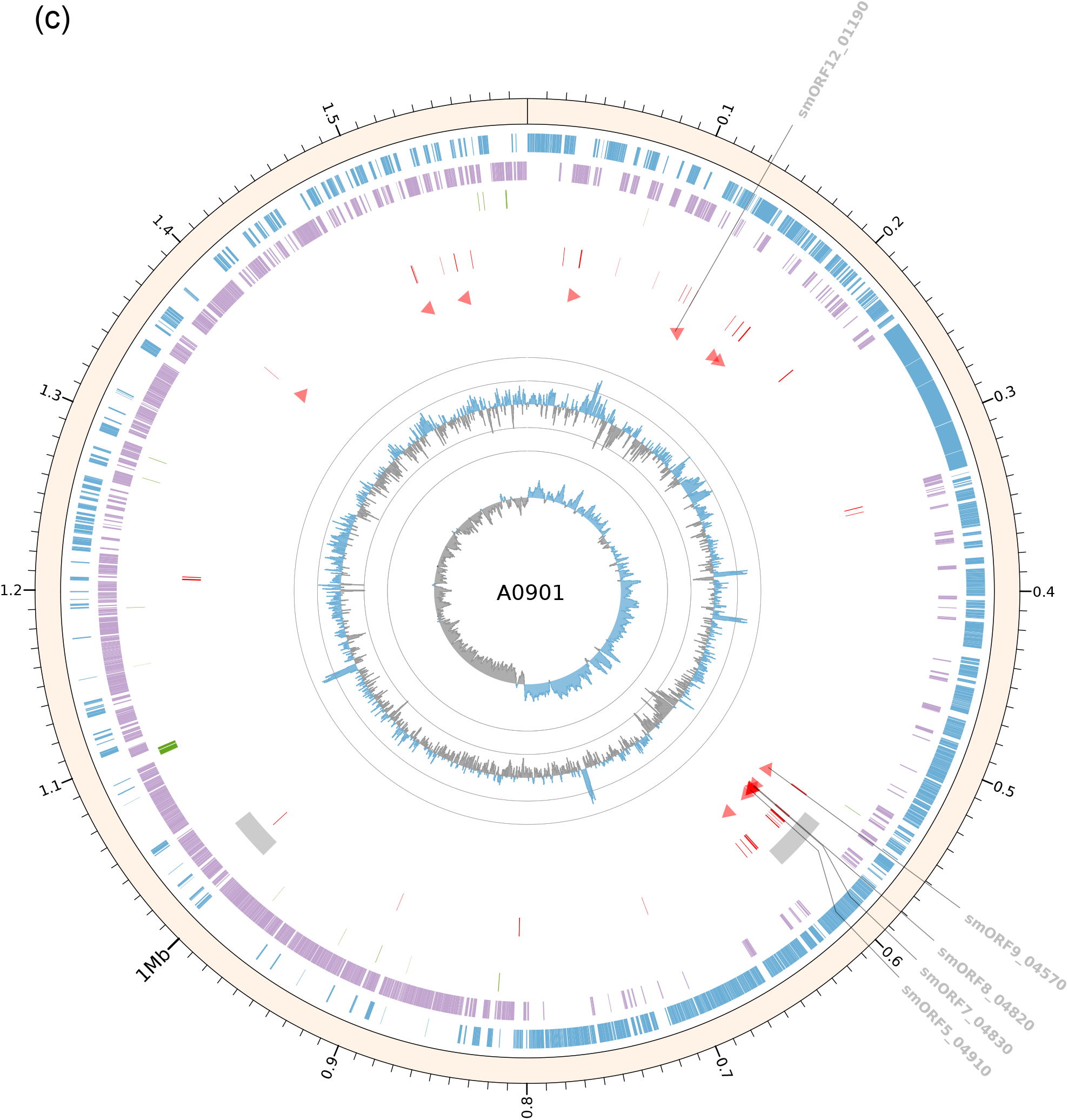
BLASTp analysis of all 1466 predicted proteins in *A. kunkeei* A0901 reveals orphan genes. (a) On X-axis, each barplot represents the number of homologs that can be found for each protein-coding gene (CDS) listed on the Y-axis. For example, the first fourteen genes on Y-axis showed zero hits on the X-axis. This suggests fourteen CDS in *A. kunkeei* A0901 had no homologs based on the analysis of BLASTp search against the NR database (excluding the self-group “*A. kunkeei*” with taxonomic ids 148814, 1419324, 1423768). Similarly, for the 1329 CDS, it showed 100 homologs. (b) Each bar plot represents the distribution of each orphan gene (on the X-axis) and the predicted protein sequence length (on the Y-axis) for the top five categories of BLASTp results from Fig S1a. The green bars and green labels on the X-axis indicate selected orphan genes to express and test in *Escherichia coli* (see, methods). (c) The complete genome of *A. kunkeei* A0901. The outer to inner circles of the circos plot (http://circos.ca) represents the genomic features of *A. kunkeei* A0901. The outer circle shows the size of the *A. kunkeei* A0901 genome. The inner circles with blue and purple blocks represent the genes that are predicted to be transcribed from the leading and lagging strands, respectively. The circle with the green blocks represents the predicted rRNA, tRNA and ncRNA genes, while the circle with two grey boxes represents the predicted phage regions. The circle with red blocks indicates 44 orphan genes (see, Fig S1b). The red triangles indicates the positioning of the selected orphan genes for further screening in *Escherichia. coli*. The GC content is represented by the blue and grey “spikes” which was calculated using a sliding window of 5000 bp. Further, the grey circles give the reference scale, which oscillates between ±20% of the mean value of 36.87% GC content. The innermost track in blue (positive) and grey (negative) shows the GC skew using a sliding window of 5000 bp.

**Figure S2.**
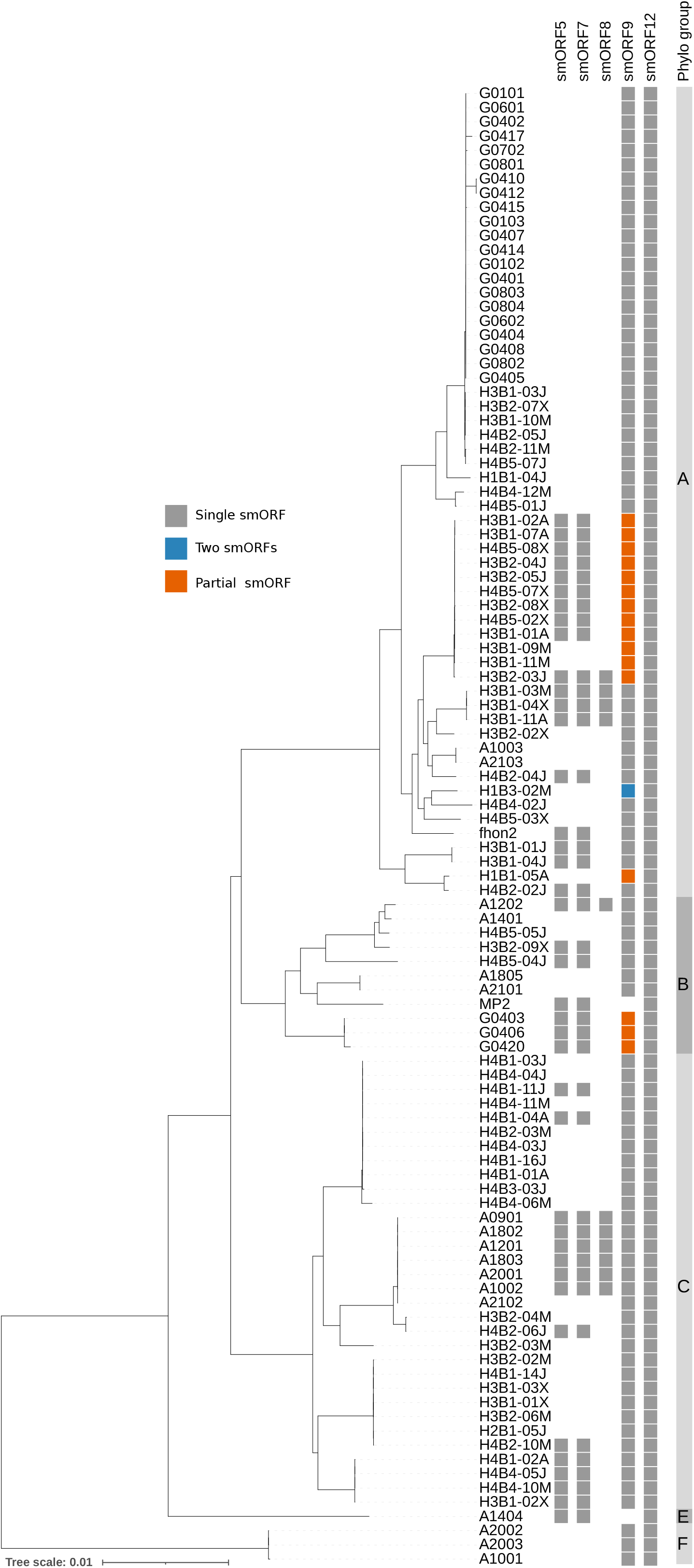
Phyletic distribution of orphan genes among different *A. kunkeei* isolates. Distribution of orphan genes among 104 *A. kunkeei* isolates. The filled boxes represent the presence of an orphan homolog whereas empty boxes indicate no sequence homolog detected in that particular species. The phylogenetic tree was taken from Dyrhage et al. (2022) *Genome Biology and Evolution* **14**, evac153.

**Figure S3.**
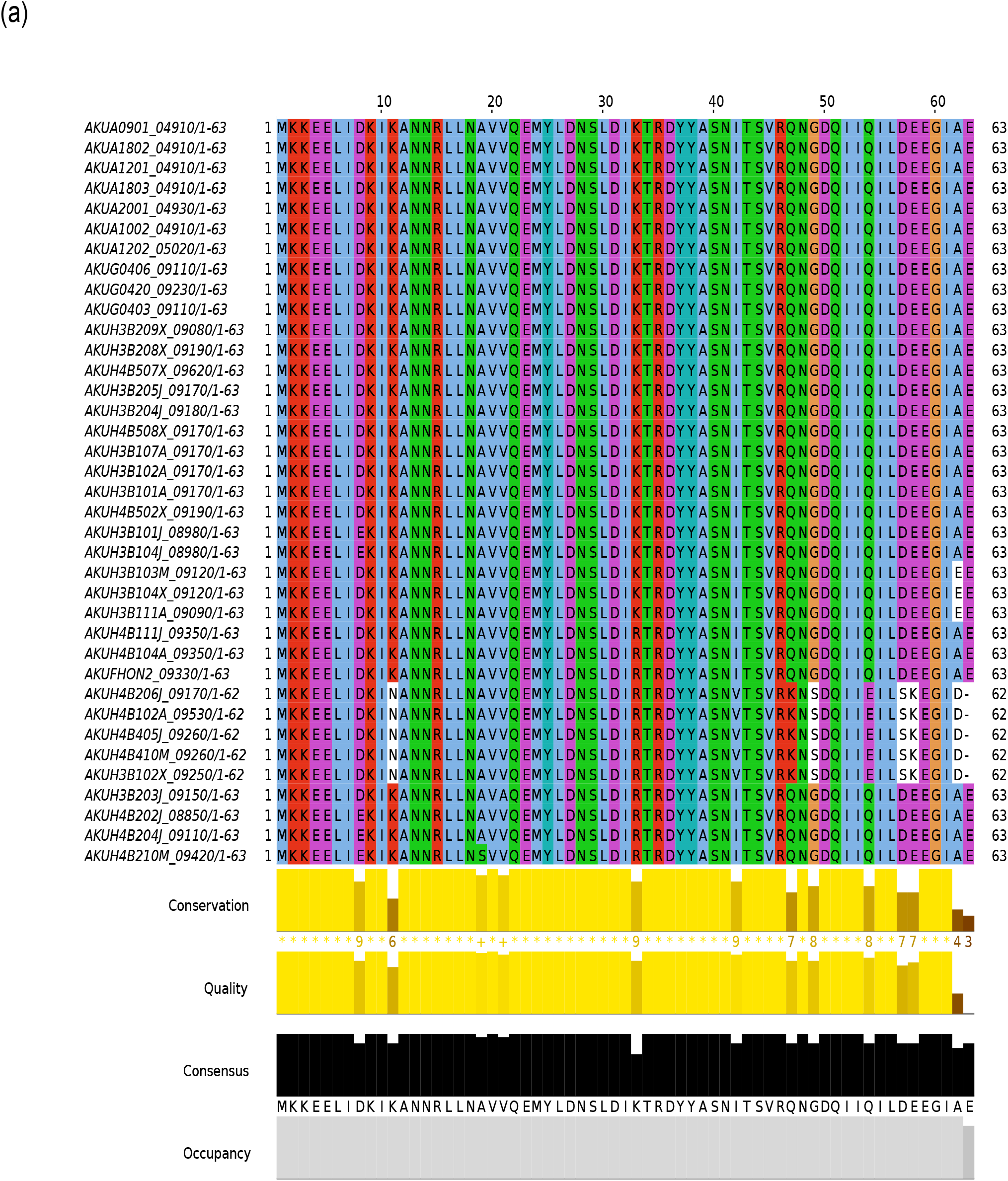
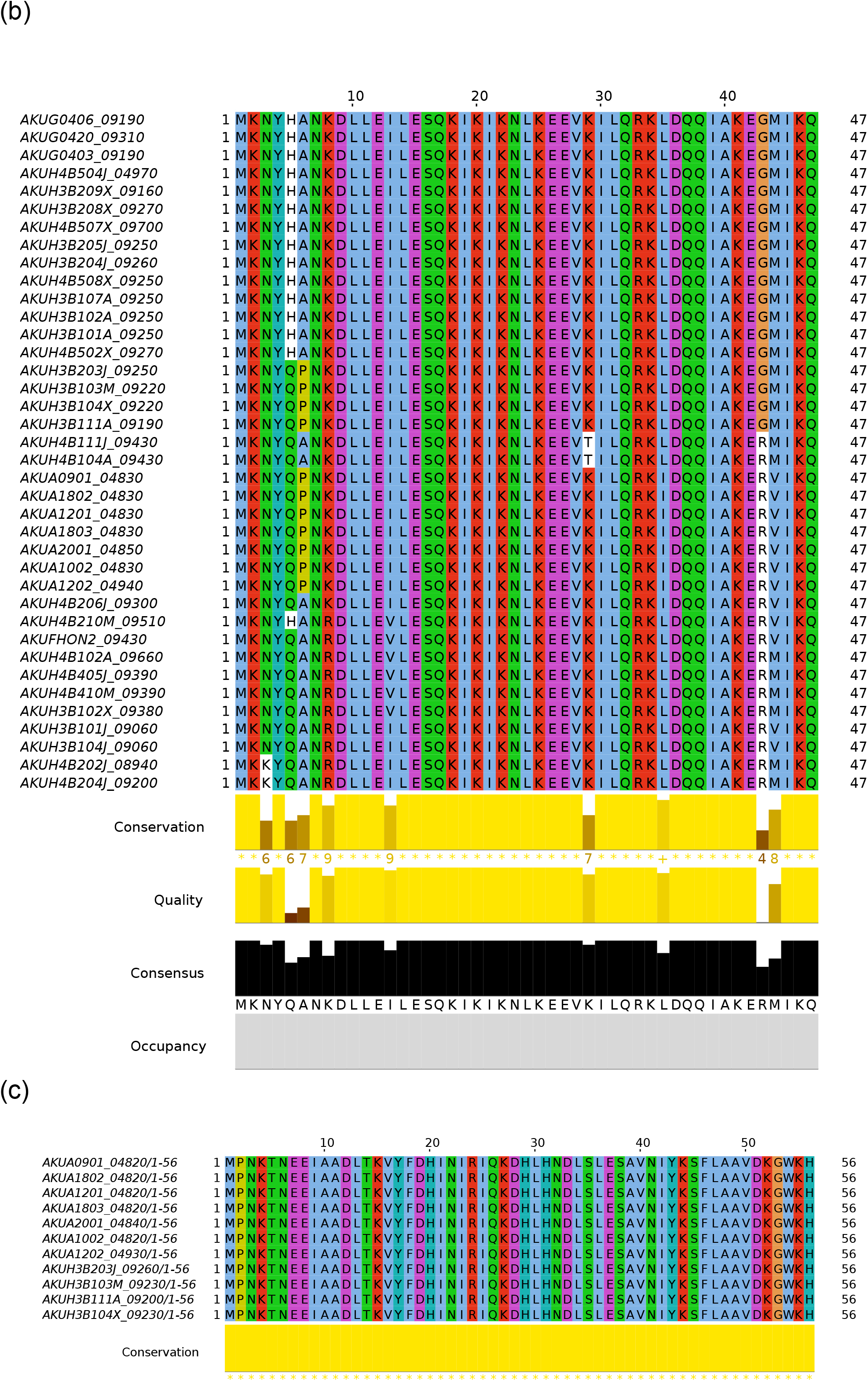
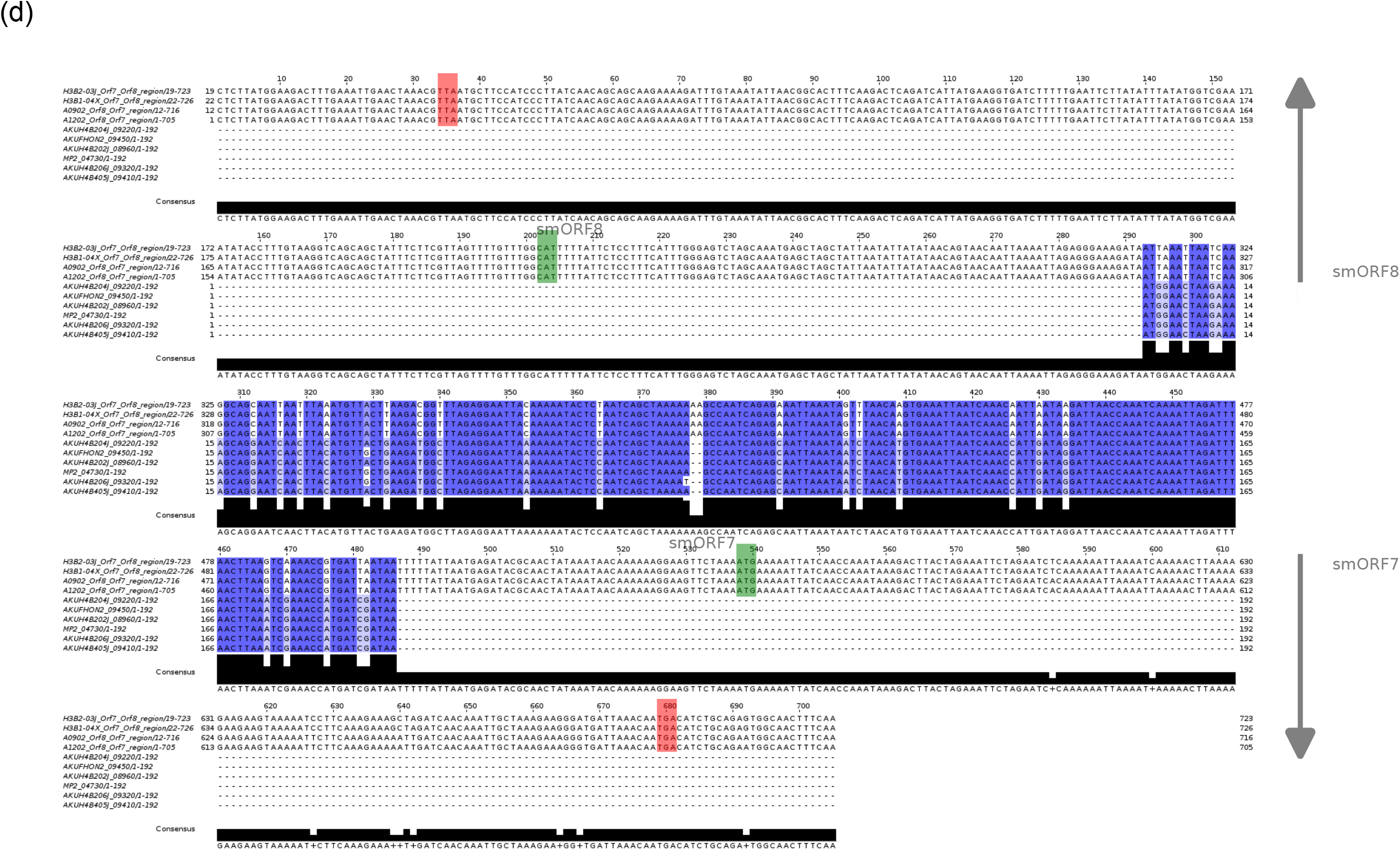
Comparison of smORF5, smORF7 and smORF8 from selected *A. kunkeei* strains. Sequence alignments of smORF protein homologs from different *A. kunkeei* strains: (a) smORF5, (b) smORF7, and (c) smORF8. (d) Sequence alignment of the smORF7 and smORF8 genes from selected *A. kunkeei* strains. The start and stop codons of the two genes are highlighted in green and red boxes, respectively.

**Figure S4.**
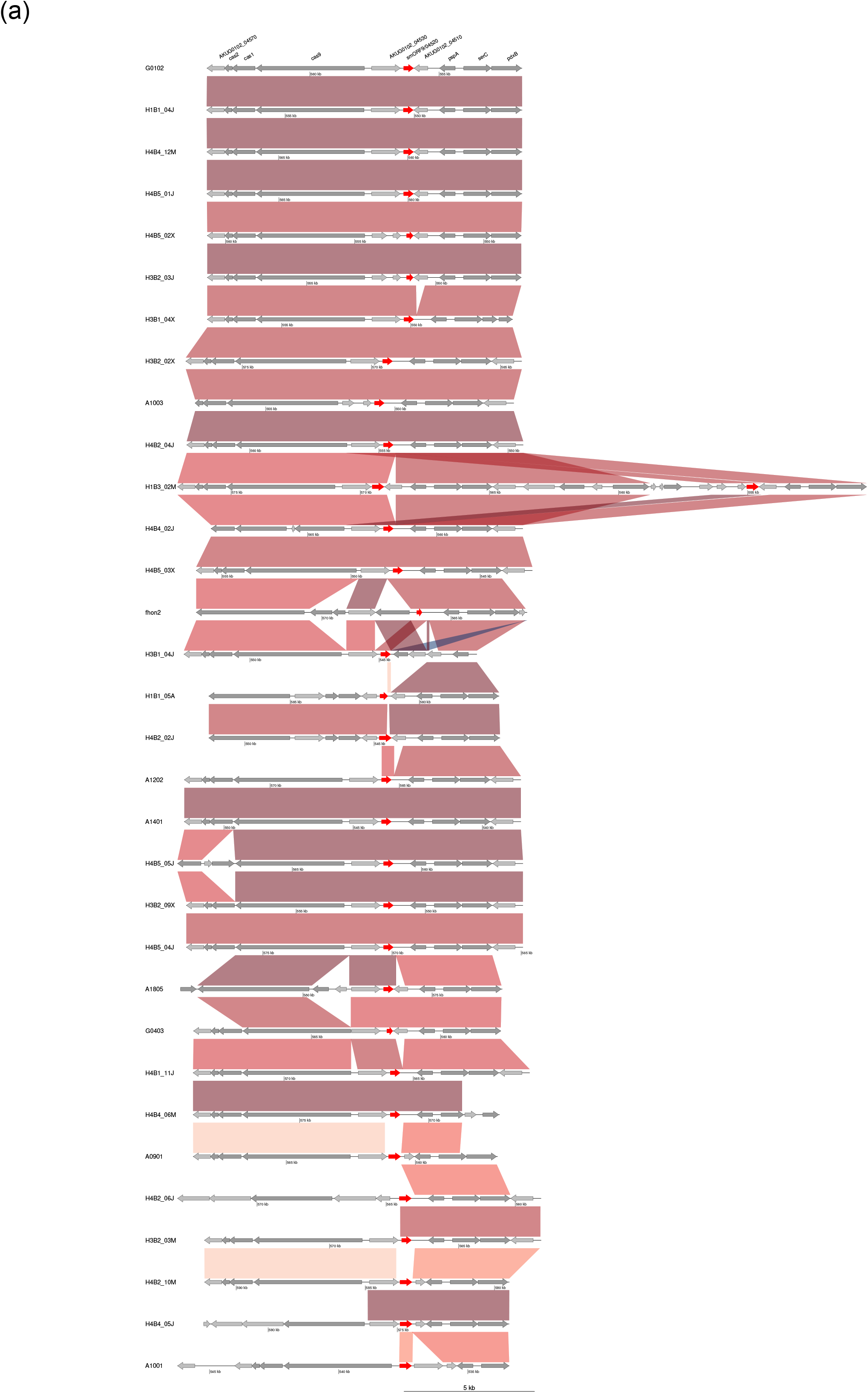
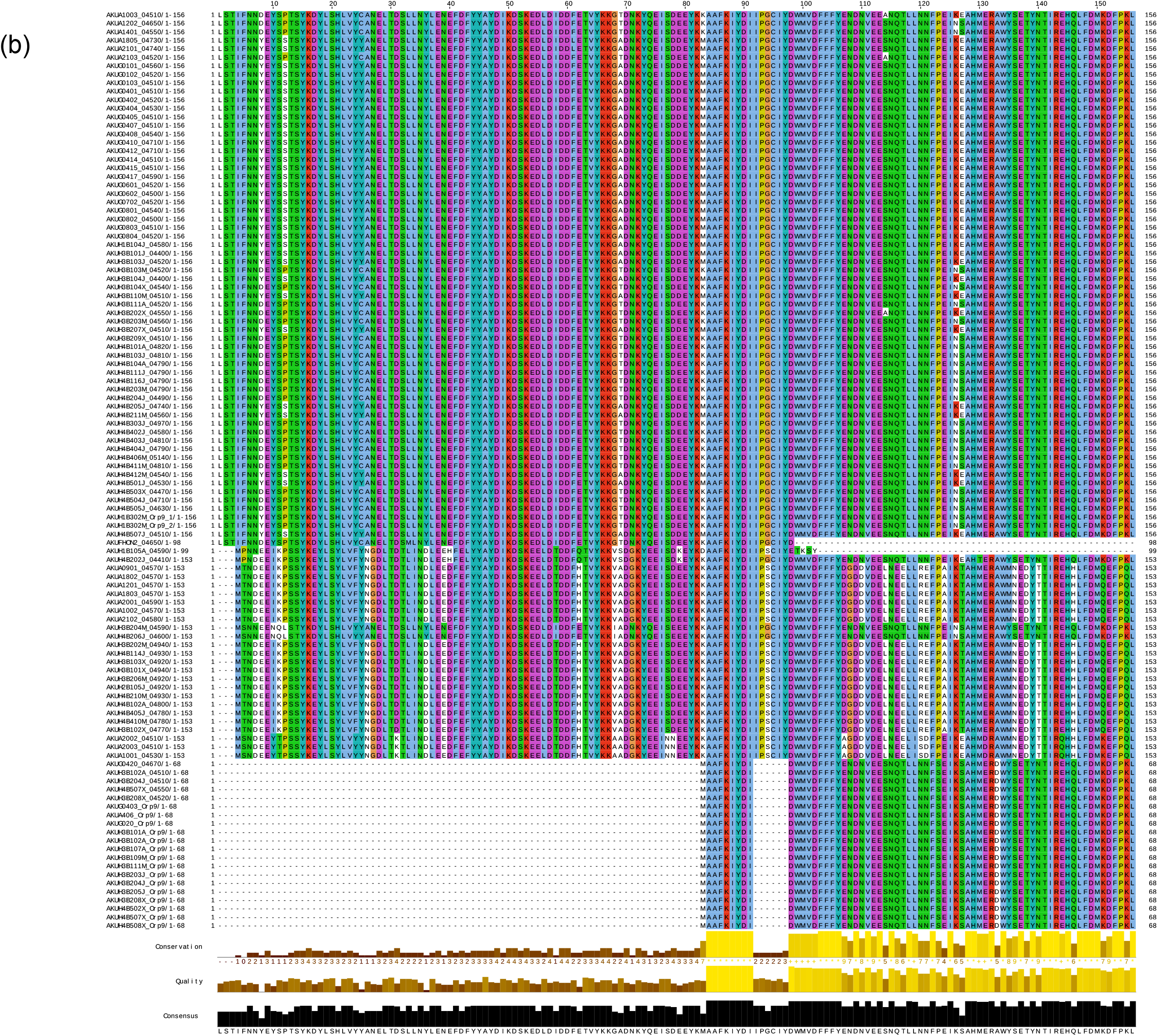
Comparison of smORF9 from selected *A. kunkeei* strains. (a) Gene synteny for the smORF9 (marked in red) gene from the selected representative members of *A. kunkeei* strains. The genes marked in grey are hypothetical protein-coding genes, whereas genes in dark grey are functionally annotated. (b) Sequence alignment for smORF9 protein homologs from *A. kunkeei* strains.

**Figure S5.**
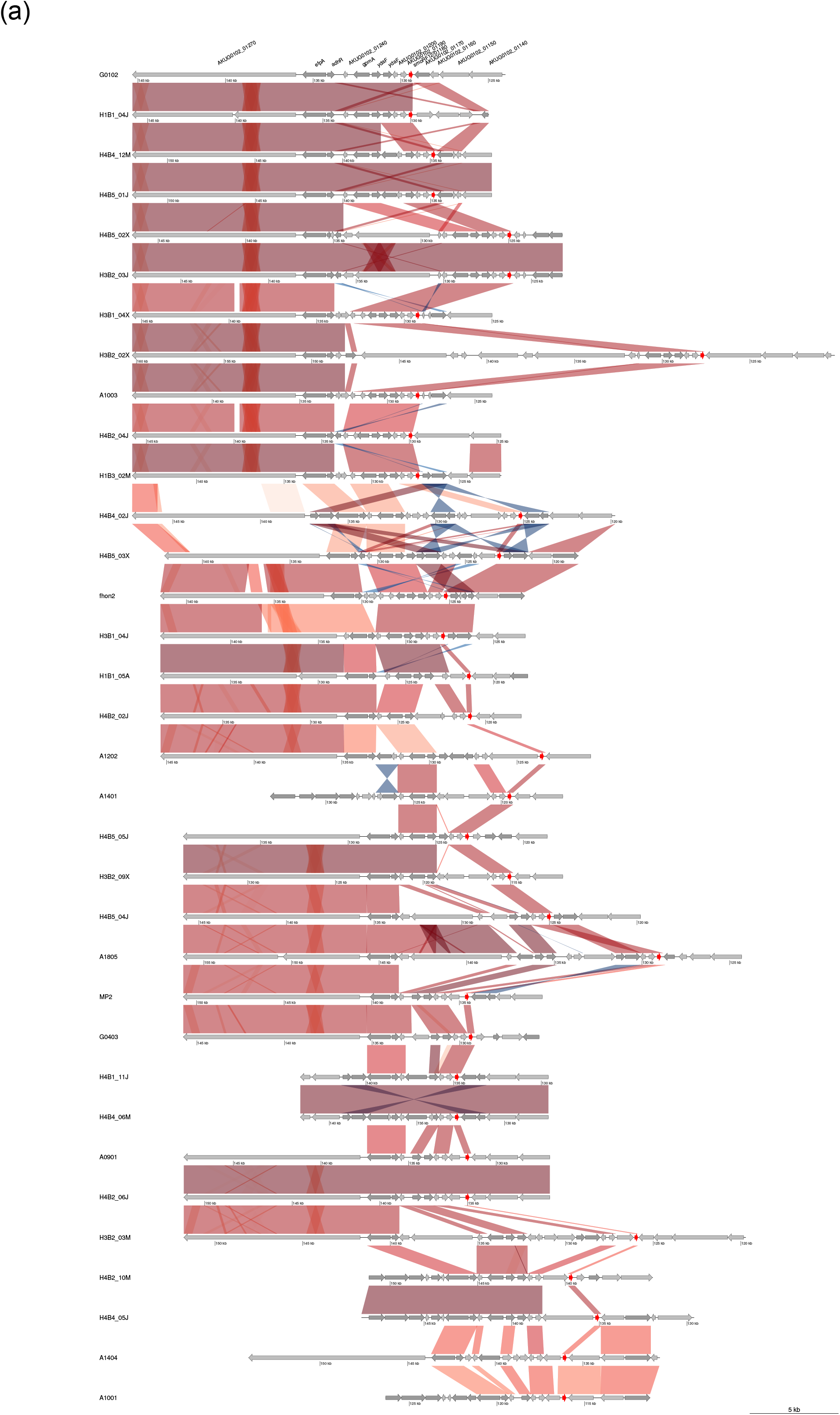
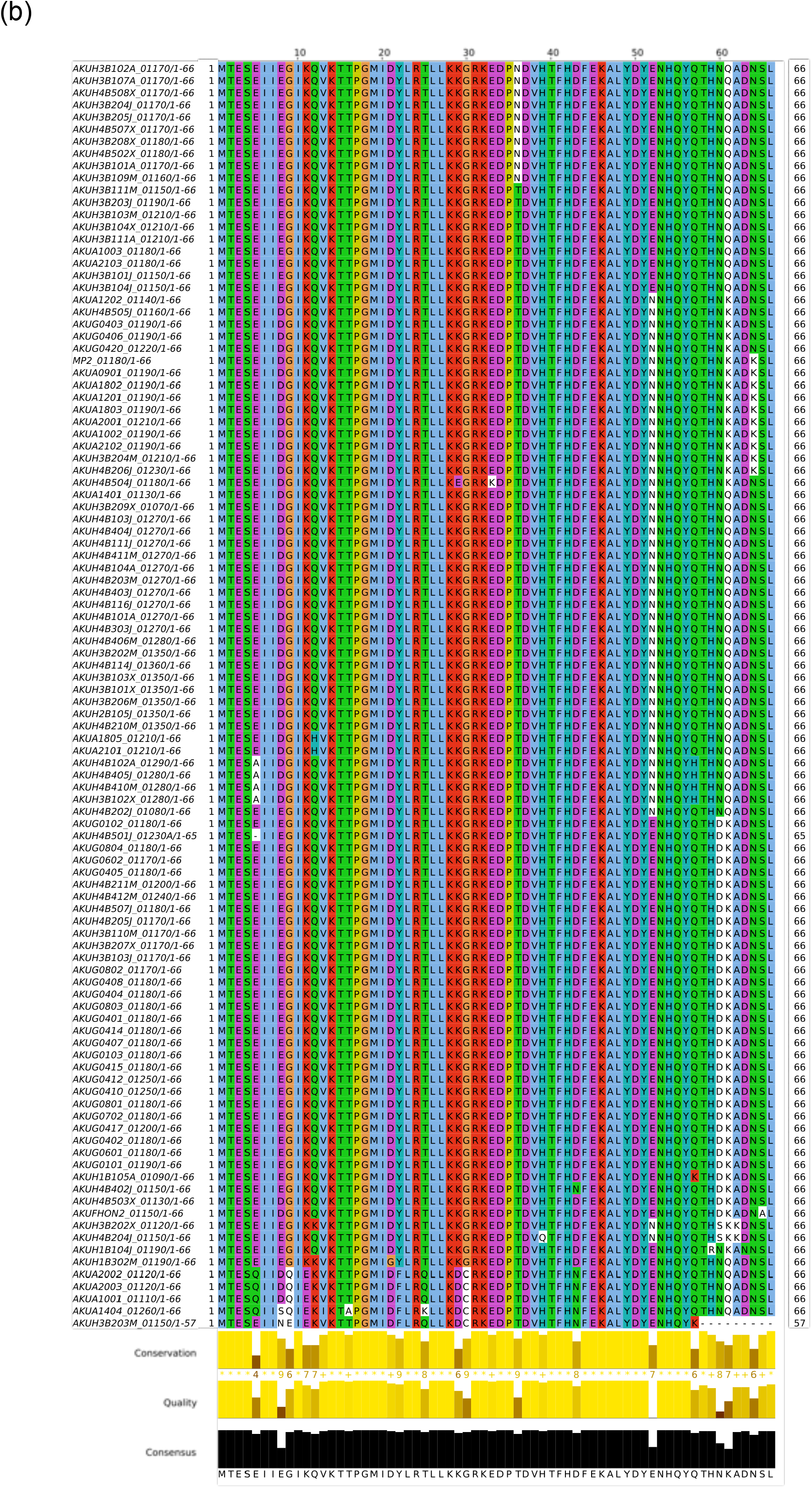
Comparison of smORF12 from the selected *A. kunkeei* strains. (a) Gene synteny for smORF12 (marked in red) gene from selected representative *A. kunkeei* strains. The genes marked in grey are hypothetical protein-coding genes, whereas genes in dark grey are functionally annotated. (b) Sequence alignment for smORF12 protein homologs from *A. kunkeei* strains.

**Figure S6.**
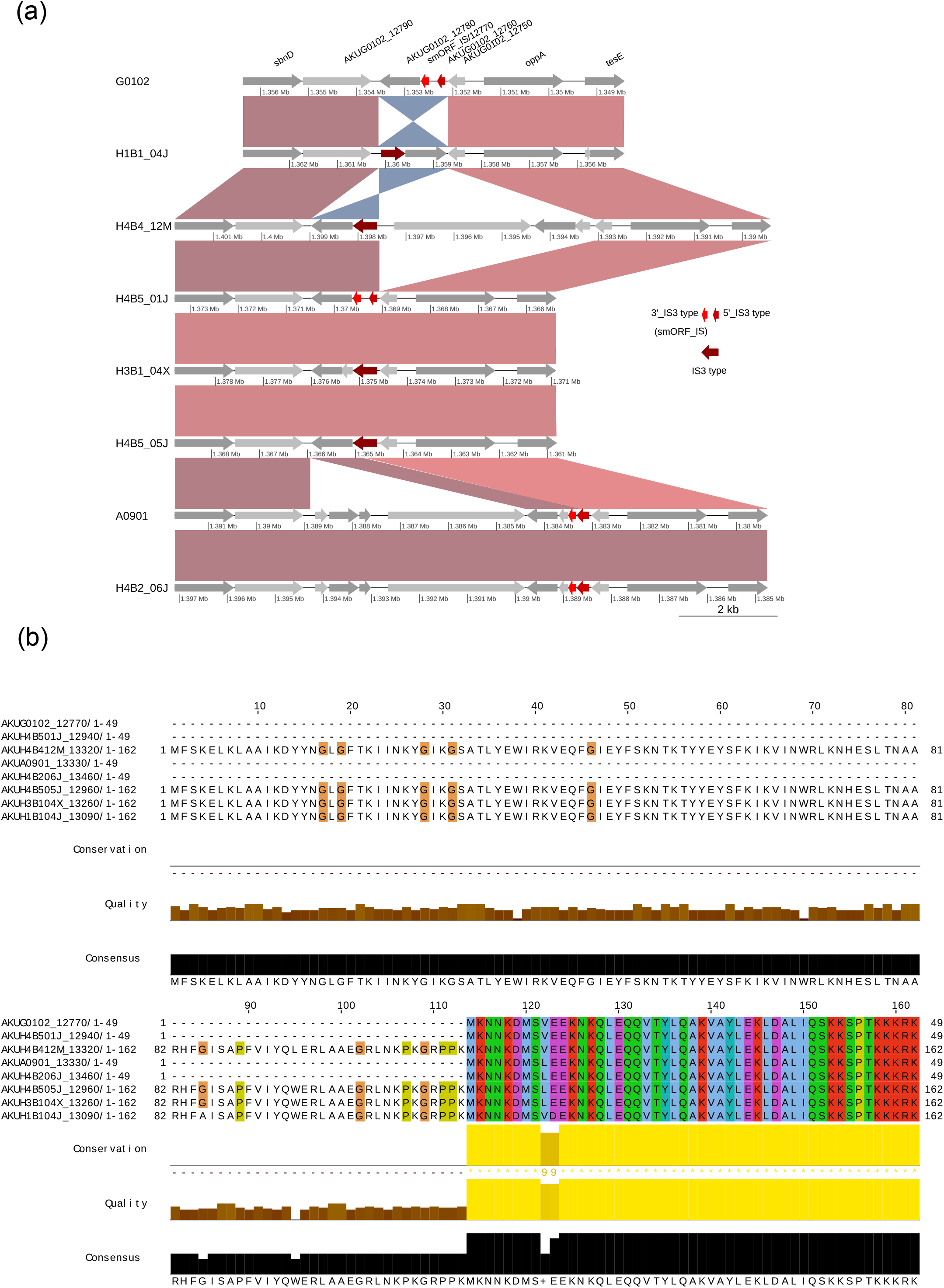
Comparison of smORF_IS from the selected *A. kunkeei* strains. (a) Gene synteny for smORF_IS (marked in red) gene from selected representative *A. kunkeei* strains. The genes marked in grey are hypothetical protein-coding genes, whereas genes in dark grey are functionally annotated. (b) Sequence alignment for smORF_IS protein homologs from *A. kunkeei* strains at the similar chromosomal location as of A0901 strain.

**Figure S7.**
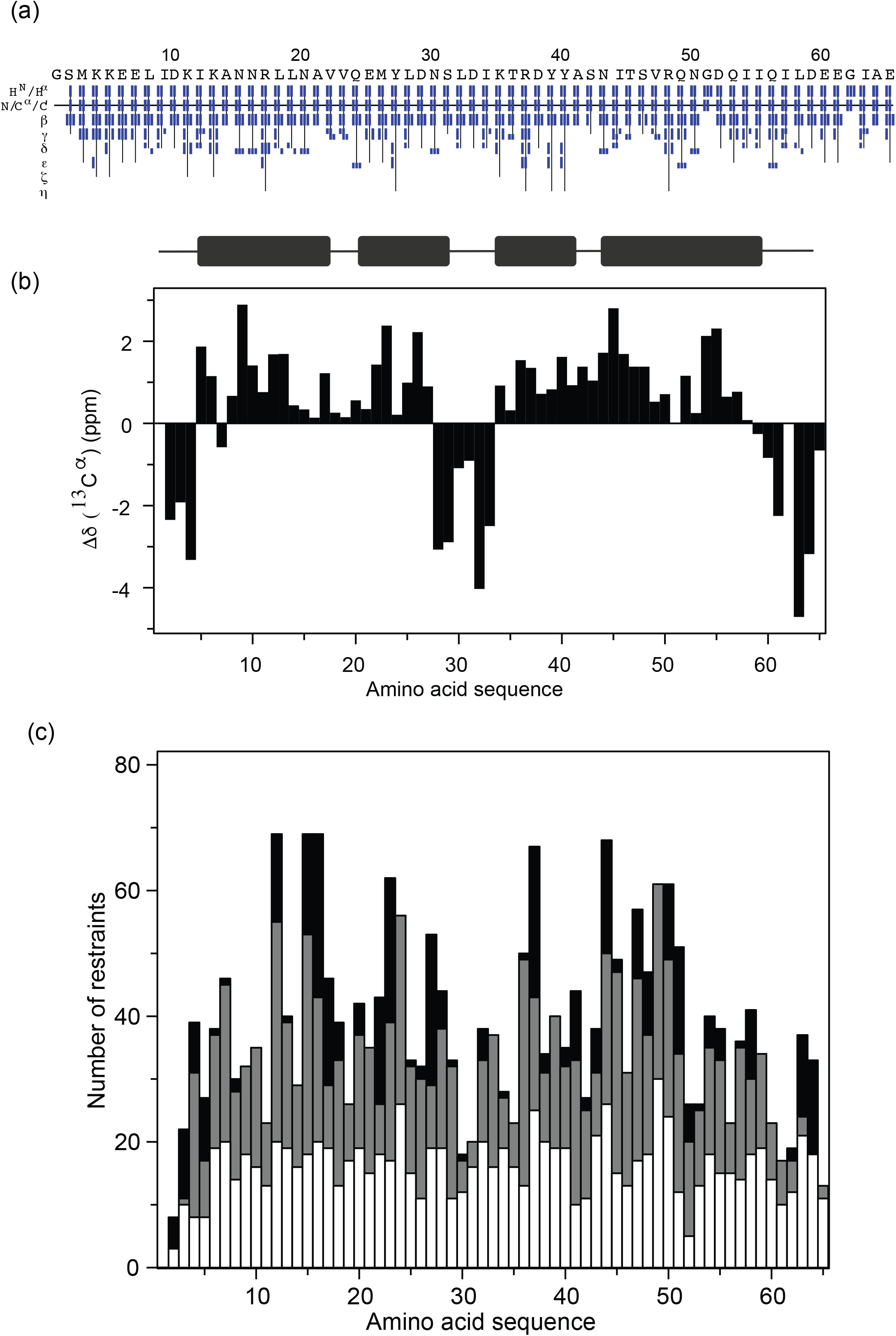
Chemical shift data and structure determination parameters for smORF5 (A0901_04910). (a) Chemical shift assignments plotted as a function of amino acid residue. The side chain atoms are indicated vertically on the left. (b) Difference of ^13^C shifts to that from random coil plotted as function of amino acid sequence. Positive values indicate helical content while negative values indicate beta-strand or random coil. c) Total number of NOE restraints per amino acid residue used for structure determination plotted as a function of the amino acid sequence. The restraints are grouped into sequential (short) (white), medium (gray) and long range (black) distances.

**Figure S8.**
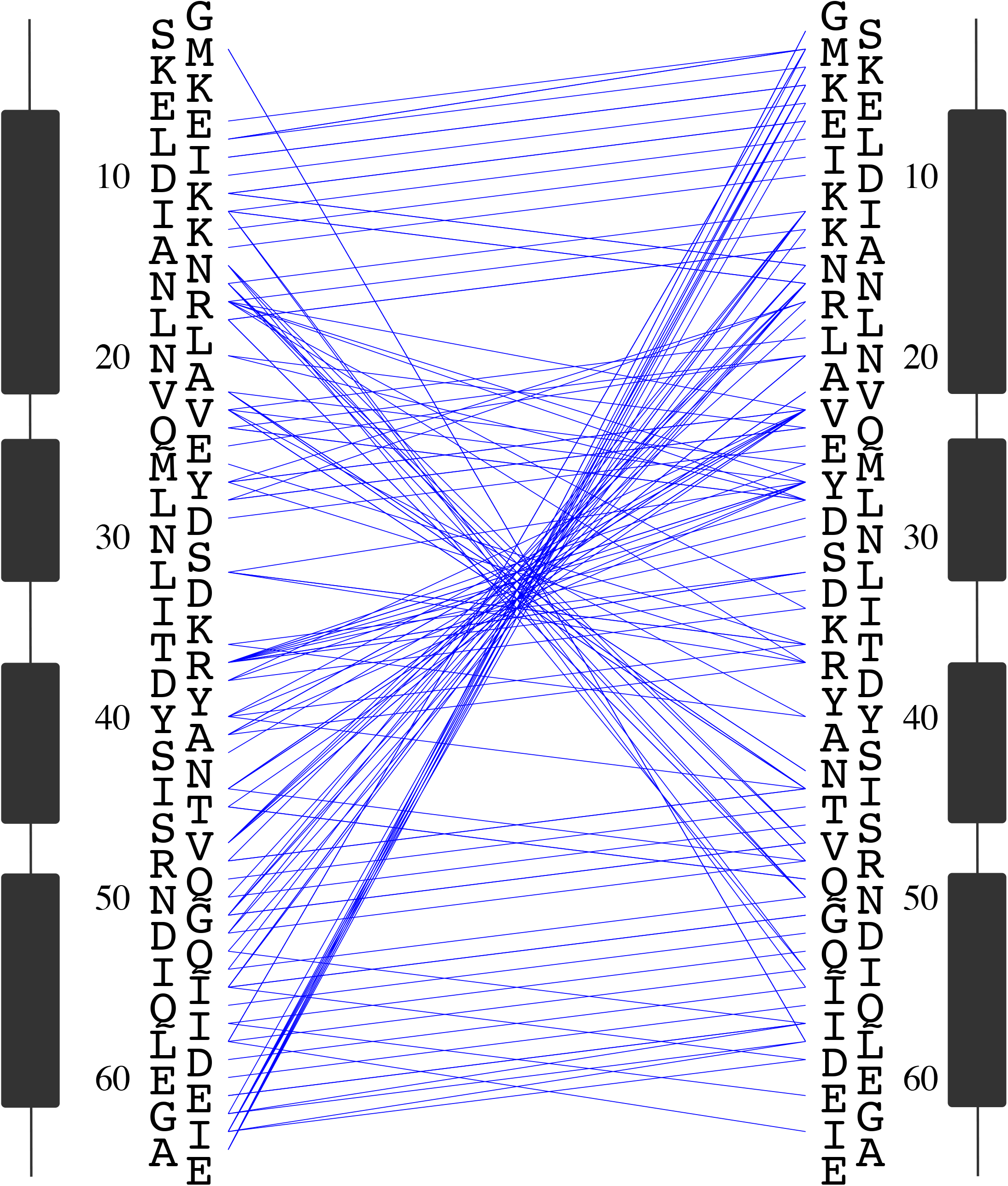
Long-range NOEs for smORF5 (A0901_04910). Each NOE is depicted by a blue line between the interacting residues. Several long-range NOEs can be seen between the N and C termini and other parts, indicating that the proposed model is well supported by the data.

**Figure S9.**
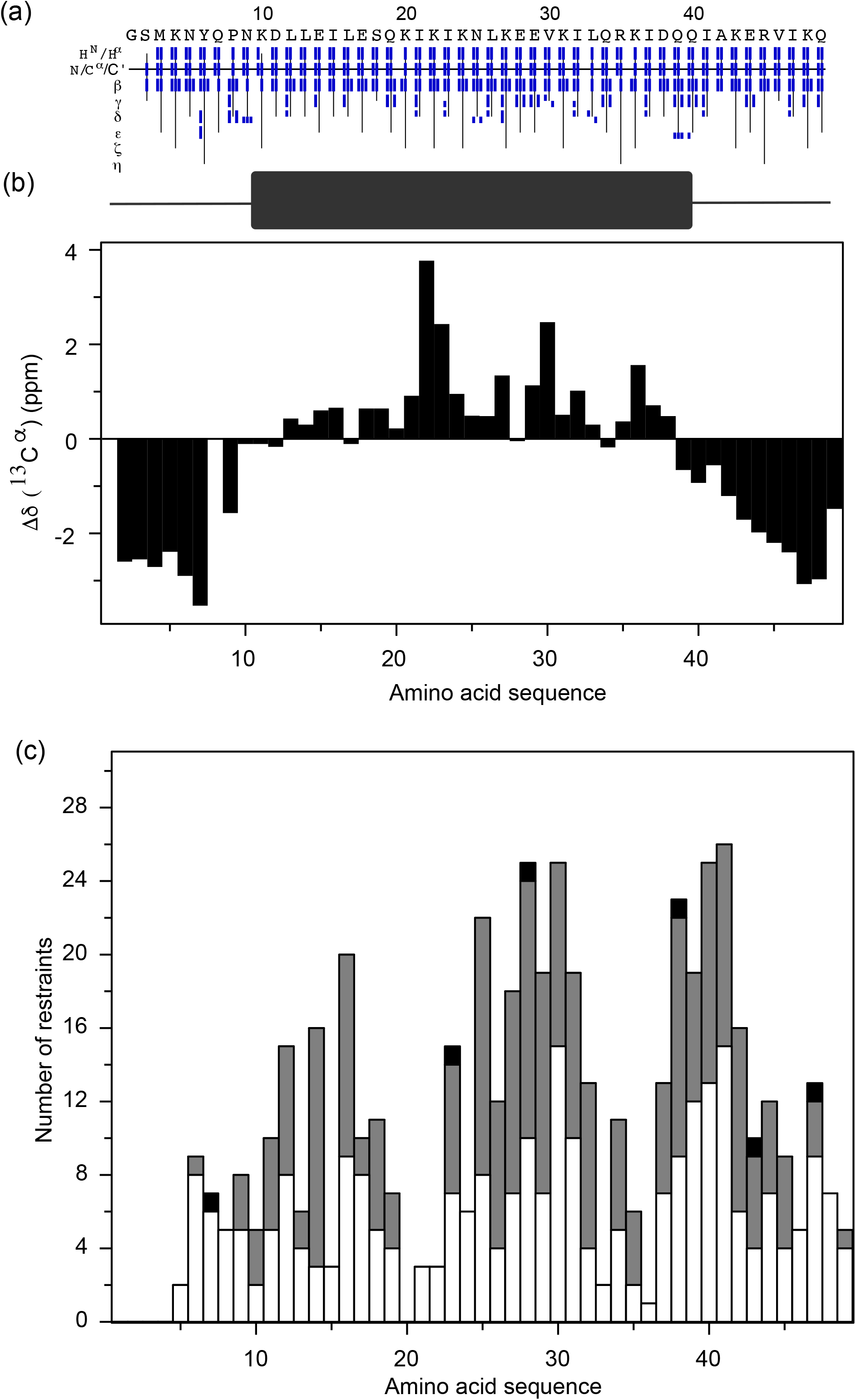
Chemical shift data and structure determination parameters for smORF7 (A0901_04830). (a) Chemical shift assignment plotted as a function of amino acid sequence. The residue atoms are indicted vertically on the left. (b) Difference of ^13^C shifts to that from random coil plotted as function of amino acid sequence. Positive values indicate helical content while negative indicate beta-strand or random coil. (c) Total number of NOE restraints per amino acid residue used for structure determination plotted as a function of the amino acid sequence. The restraints are grouped into sequential (short) (white), medium (gray) and long range (black) distances.

**Figure S10.**
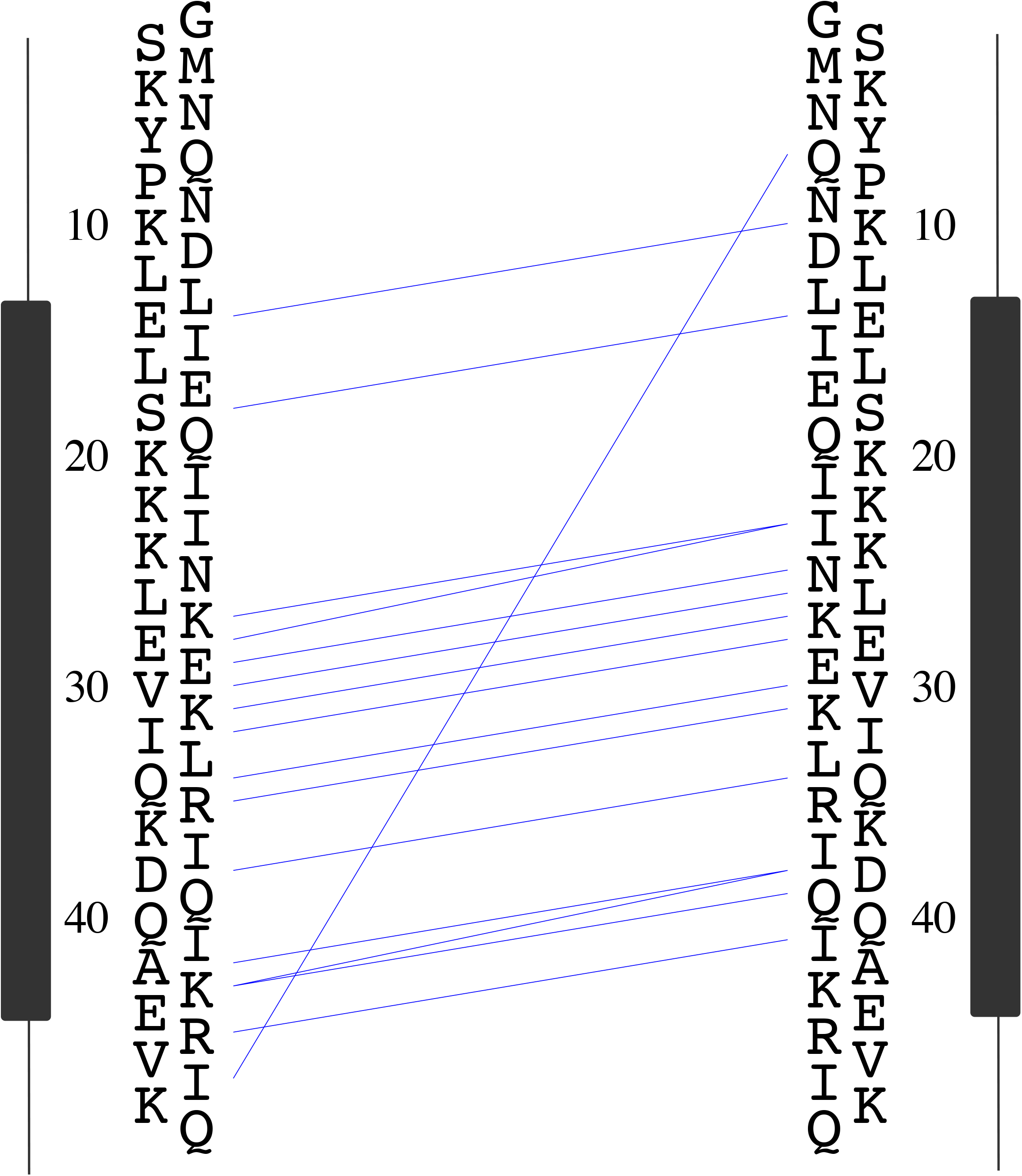
Long-range NOEs for smORF7 (A0901_04830). Each NOE is depicted by a blue line between the interacting residues. Only a single long-range and many short-range NOEs are seen, which is consistent with a fold of this nature.

**Figure S11.**
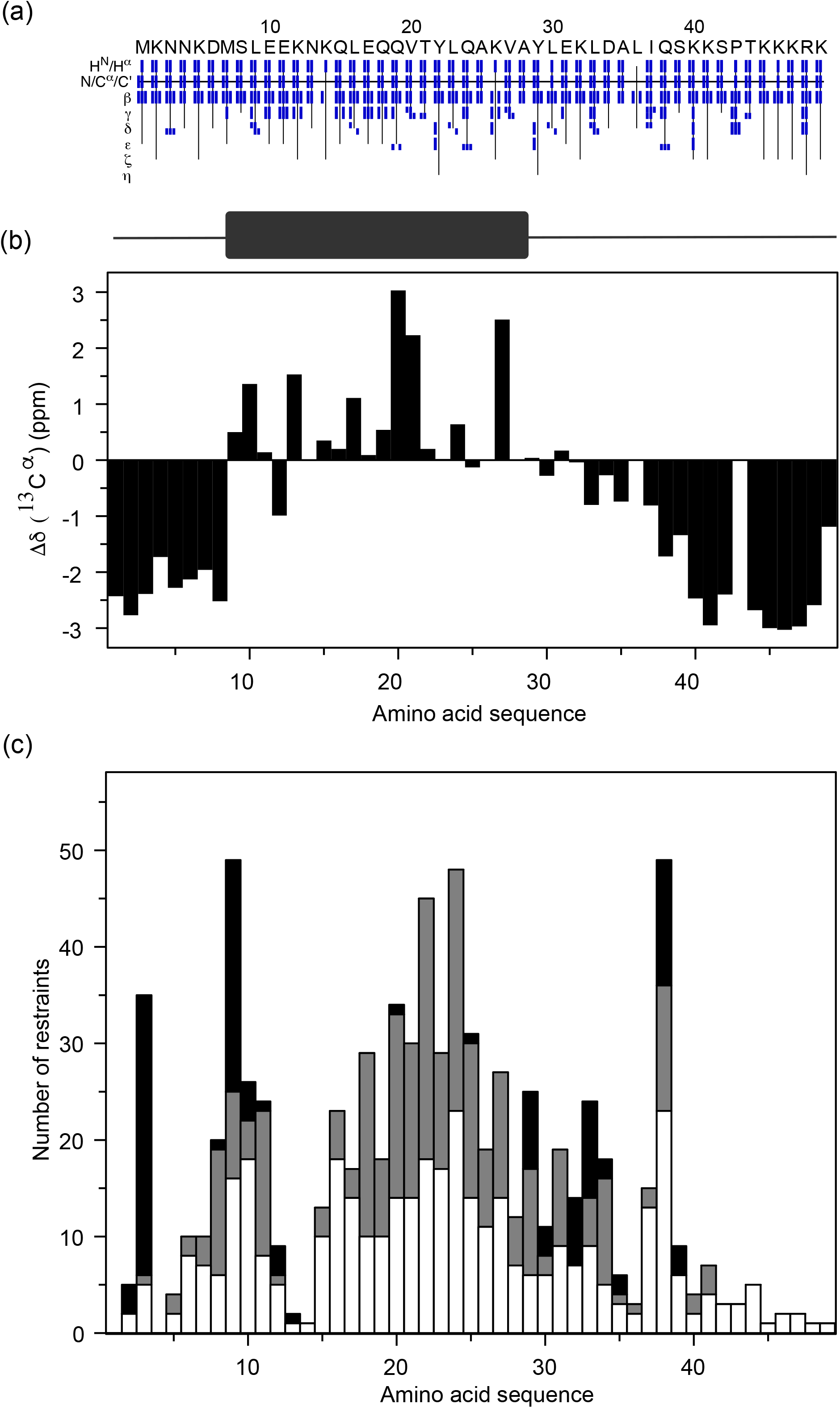
Chemical shift data and structure determination parameters for smORF_IS (A0901_13330). Chemical shift assignments plotted as a function of amino acid residue. The side chain atoms are indicated vertically on the left. (b) Difference of ^13^C shifts to that from random coil plotted as function of amino acid sequence. Positive values indicate helical content while negative indicate beta-strand or random coil. (c) Total number of NOE restraints per amino acid residue used for structure determination plotted as a function of the amino acid sequence. The restraints are grouped into sequential (short) (white), medium (gray) and long range (black) distances.

**Figure S12.**
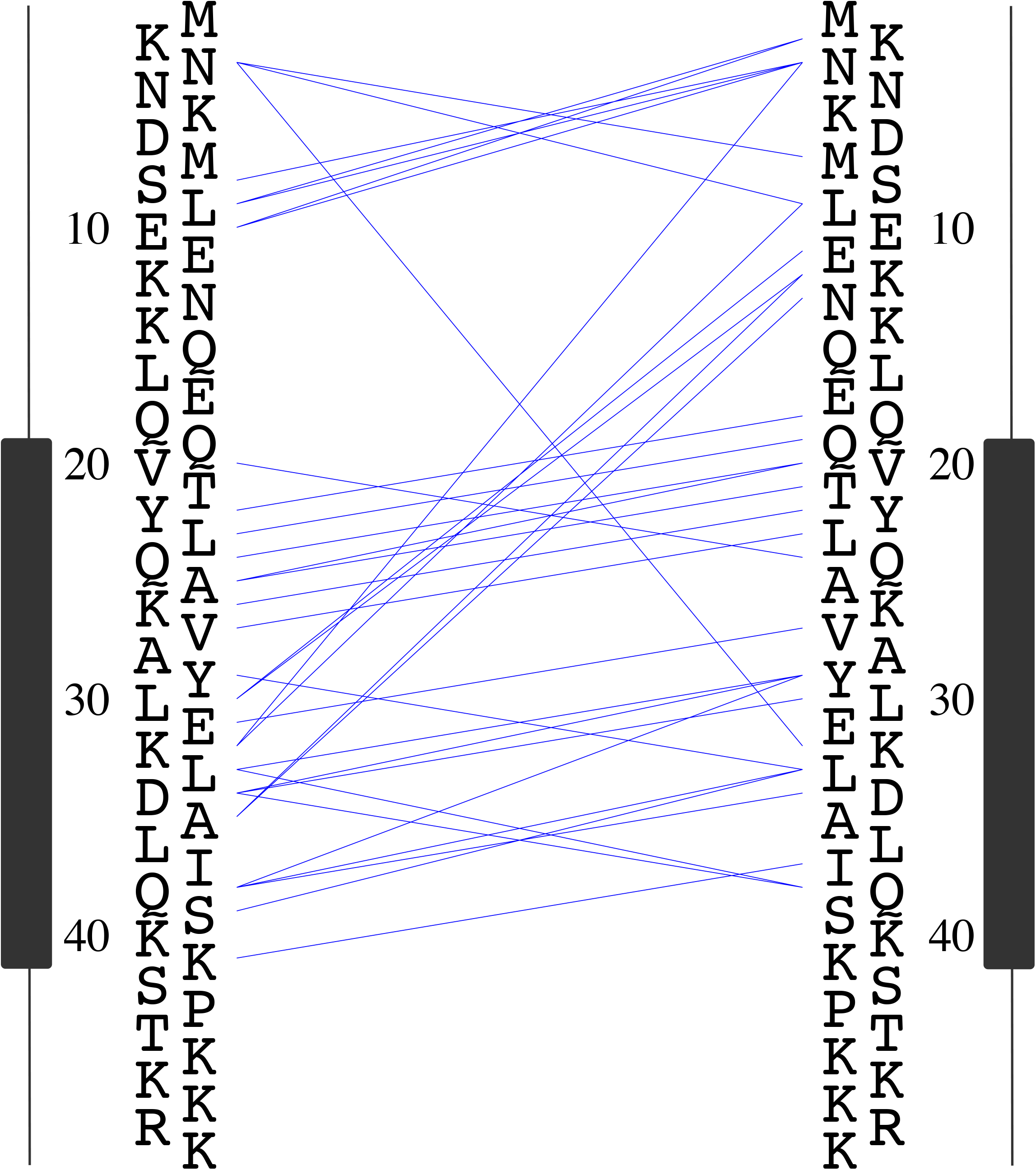
Long-range NOEs plotted from residue to residue for smORF_IS (A0901_13330). Each NOE is depicted by a blue line between the interacting residues. Despite forming only a single helix and long unstructured N and C termini, smORF_IS contains seven long-range NOEs. This means that the folded helix makes transient contacts with the unstructured region in a slow exchange (μs-ms time regime).

### Supplementary Data

**Table S1:** Compilation of top five BLASTp hits summary categories from *Apilactobacillus kunkeei* A0901 strain based on the NCBI nr-database (ver 2023-01-10). All BLASTp results were obtained by performing against the NCBI database by excluding the self-group “*Apilactobacillus kunkeei*” (Taxonomy ids: 148814, 1419324, 1423768) with e-value 1e-03 along with the default settings.

**Table S2a:** Compilation of top five categories species summary based on Table S1.

**Table S2b:** Average nucleotide identity comparison for the species closely related to the genus Apilactobacillus (Table S2a). Pairwise ANI analysis is performed using the tool fastANI ver1.33

**Table S3:** List of smORFs detected in proteomics dataset in A0901 strains during the growth and increasing concentrations of CPX amount (see, methods).

**Table S4:** List of smORFs in numerical order and with identifier.

**Table S5:** Raw data, and fitted and calculated parameters from **Fig. 3**.

